# Ventral frontostriatal circuitry mediates the computation of reinforcement from symbolic gains and losses

**DOI:** 10.1101/2024.04.03.587097

**Authors:** Hua Tang, Ramon Bartolo, Bruno B. Averbeck

## Abstract

Reinforcement learning (RL), particularly in primates, is often driven by symbolic outcomes. However, it is usually studied with primary reinforcers. To examine the neural mechanisms underlying learning from symbolic outcomes, we trained monkeys on a task in which they learned to choose options that led to gains of tokens and avoid choosing options that led to losses of tokens. We then recorded simultaneously from the orbitofrontal cortex (OFC), ventral striatum (VS), amygdala (AMY), and the mediodorsal thalamus (MDt). We found that the OFC played a dominant role in coding token outcomes and token prediction errors. The other areas contributed complementary functions with the VS coding appetitive outcomes and the AMY coding the salience of outcomes. The MDt coded actions and relayed information about tokens between the OFC and VS. Thus, OFC leads the process of symbolic reinforcement learning in the ventral frontostriatal circuitry.

## INTRODUCTION

Reinforcement learning (RL) is an adaptive process by which agents learn to make choices to gain rewards over some future time horizon^1^. These processes are often studied in animal models using tasks in which choices lead to primary rewards^2,3^. However, in many situations, choices lead to symbolic outcomes that lead to rewards in the future. Humans are motivated by and will work for symbolic outcomes in the form of money^4,5^. Animals also readily learn to make choices that lead to symbolic reinforcers, often in the form of tokens. For example, numerous studies have shown that primates will learn to make decisions to maximize the accumulation of tokens that are periodically converted to primary reinforcers^6-9^.

Early studies established an important role for the striatum and the dopamine innervation of the striatum in RL^10-13^. Subsequent work has shown that additional structures, including a network composed of the OFC^14,15^, AMY^16,17^, and MDt^18^ are also involved in RL^19^. Although causal manipulations and neurophysiology have shown that each of these areas plays a role in RL, the results across tasks and areas are not always consistent. For example, lesions to the VS cause deficits in tasks that use probabilistic delivery of primary reinforcers^17^. However, these deficits are limited to learning to associate rewards with objects and not actions^20^. This is not consistent with actor-critic models of RL that suggest that VS stores a general state value representation for policy learning^21,22^. Similarly, AMY and VS appear to play stronger roles in probabilistic bandit tasks but not in token-based RL tasks, in which tokens are used as reinforcers^1,7,23^. In token-based RL, VS is only involved in learning to discriminate between relative gains and plays no role in learning to discriminate between gains and losses^7^. The AMY, on the other hand, appears to play a limited role in token-based RL^23^. Previous neurophysiology studies have shown that token-based learning may engage cortical networks more than subcortical networks^8,9^, whereas learning based directly on primary rewards may preferentially engage subcortical networks^24,25^. Beyond these examples, some learning-related behaviors may rely more on inference processes than incremental value updates that characterize RL and may, therefore, tap into different networks^26,27^. Thus, the way in which the ventral cortico-striatal-thalamo-cortical network, including the OFC, VS, AMY, and MDt, orchestrates RL, particularly with symbolic reinforcers, is unclear.

In the present study, we carried out simultaneous neurophysiology recordings across the ventral cortico-striatal-thalamo-cortical network using a token-based RL task. We examined single neuron, population, and network computations underlying the performance of the task. The results show that the OFC played a dominant role in the computations relevant to learning from symbolic reinforcers. Token outcomes were coded earlier and more strongly in OFC. Conversely, the VS was characterized by specific and unique coding of appetitive choices and outcomes. The AMY was characterized by coding of salience, a finding not apparent in previous work that used only probabilistic appetitive outcomes. The MDt did not appear to contribute a unique computation. However, it played an important role in mediating the interaction of OFC and VS during the calculation of token outcomes. Together, these results define the unique and shared contributions of the ventral frontostriatal circuitry to learning from symbolic reinforcers.

## RESULTS

Two rhesus monkeys were trained on a two-armed bandit task. In this task, the monkeys collected tokens, which were periodically exchanged for juice rewards (Figures 1A-B). In every block of 108 trials, we introduced four novel images. In each trial, two of the four images were presented on the screen. The choice of an image led stochastically to one of -2, -1, +1, or +2 tokens (Figure 1B). The image-outcome relationships were unknown to the monkeys at the start of the block. Monkeys had to learn the values of the images by choosing one of them and observing the outcome. Token outcomes were stochastic such that in 75% of the trials, the monkeys received the number of tokens associated with the chosen image, and in 25% of the trials, the number of tokens did not change. Tokens were accumulated across trials and were cashed out every four to six trials, with one drop of juice for each token.

**Figure 1.**
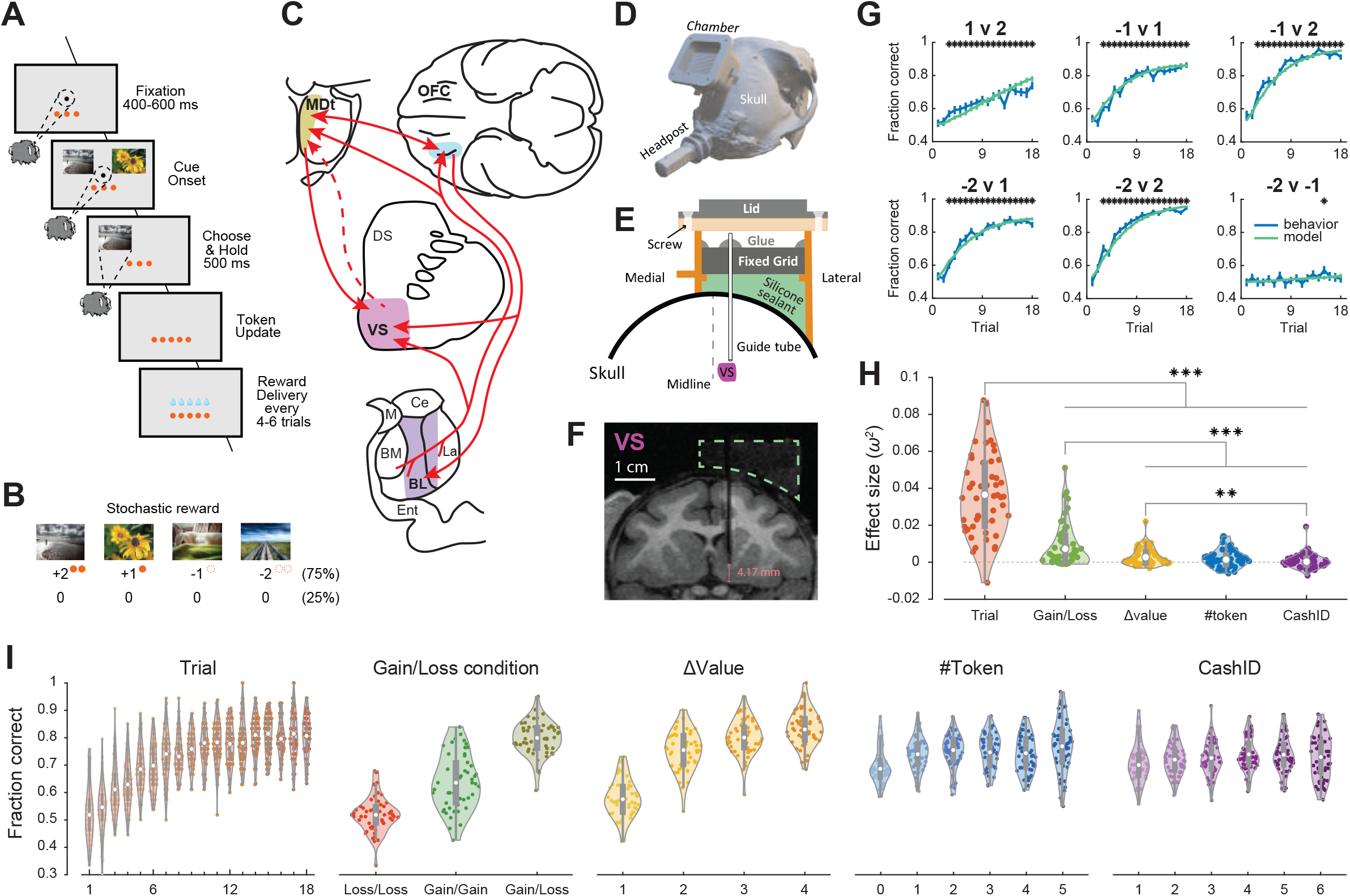
Task, behavior, recording method, and recorded areas. (A-B) Behavioral task. (A) Structure of an individual trial. Successive frames illustrate the sequence of events. In each trial, only two of the four images were presented. Monkeys chose between them and gained or lost (stochastic: 75% change, 25% no change) the corresponding number of tokens. Choices could be made as soon as the images were shown. Accumulated tokens were cashed out every four to six trials, with one drop of juice for each token. (B) In each block, the monkeys learned the values (+2, +1, −1, −2) of four novel images. (C) Schematic recording areas, including the orbitofrontal cortex (OFC), ventral striatum (VS), basolateral amygdala (AMY), and mediodorsal thalamus (MDt). Red arrows indicate the anatomical connections between these areas. The dashed line indicates an indirect connection. (D-F) Semi-chronic recording. (D) A chamber was implanted. (E) Then, guide tubes were inserted into each target area through a grid. The guide tubes stopped above the target areas and stayed in the brain across recording days. The empty space in the chamber was filled with silicone sealant to prevent potential infections. Multi-site linear probes were lowered into target areas through the guide tubes every recording day. (F) A coronal section view of the chamber shows the shadow of a guide tube and the target area under MRI. The dashed green polygon indicates the silicone sealant. (G-I) Choice behavior. (G) Fraction of choosing the image with a higher value in each condition. Blue lines represent monkey behavior, and green lines represent the prediction of the reinforcement learning model. Error bars represent mean ± SEM. The black asterisks indicate a significant difference in monkey behavior from the change level (one-sample t-test, p < 0.01). (H) The effect size of each task variable. Paired t-test, **p < 0.01, ***p < 0.001. Non-significant pairs are not indicated. (I) Choice behavior as a function of the task variables. Results were averaged from two monkeys, n = 50 sessions. Violin plots showing the range of values calculated across all sessions (center dot, mean; box, 25th to 75th percentiles; whiskers, ± 1.5 × the interquartile range; dots, each session; shade, density curve).

The primary reward was apple juice (#juice), but it was only delivered every four to six trials. In each trial, the monkeys had a chance to gain or lose tokens by choosing one of the images. The chosen value information was first carried by the images (cValue), which predicted the change of tokens (Δtoken). The number of accumulated tokens (#token) was always presented on the screen. So, value information appeared in different forms, including cValue, Δtoken, #token, and #juice (Figure S1A). The following analyses focus on the neural representation and interactions of these forms of value.

### Choice behavior was influenced by gaining/losing tokens and token numbers

We quantified choice behavior by measuring the fraction of times the monkeys chose the image associated with a higher value in each condition (e.g., choose +1 when presented with -1 and +1 images). The monkeys learned to distinguish the values of two images shown on the screen within about 10 trials for the Gain/Loss and Gain/Gain conditions but learned minimally in the Loss/Loss condition (Figure 1G and Figure S1B). We also fit a Rescorla-Wagner (RW) reinforcement learning model to the choice behavior (Figure 1G), which has been explored previously^7^. The model is used below to examine token reward prediction error (RPE). To quantify how the choice behavior (i.e., whether they chose the better option) was affected by task variables, we fit a multi-way ANOVA model to it (Figures 1H-I). The choice behavior was significantly modulated by the number of observations (Trial, *F*_17, 44950_ = 109.98, p < 0.001), Gain/Loss condition (Gain/Loss, *F*_1, 44950_ = 286.91, p < 0.001), value difference of two options (Δvalue, *F*_3, 44950_ = 73.37, p < 0.001), and the number of tokens before choice (#token, *F*_10, 44950_ = 8.68, p < 0.001), but not by the number of trials since last token cashout (CashID, *F*_5, 44950_ = 2.07, p = 0.066). The number of observations indicates the learning process, both the Gain/Loss condition and Δvalue indicate the updating of tokens, #token indicates the accumulated value, and the CashID indicates the probability of receiving a primary reward. The result indicates that the monkeys understood the meaning of tokens and adjusted their behavior to get more tokens as learning progressed.

### Neural encoding of choices, primary and symbolic reinforcers

We performed simultaneous recordings of population neural activity across four regions of the ventral cortico-striatal-thalamo-cortical network (Figure 1C and Figures S1C-E) from two macaque monkeys using multi-site linear probes. We collected 606 neurons in the 13L region of the orbitofrontal cortex (OFC), 829 neurons in the core region of the ventral striatum (VS), 1607 neurons in the basolateral amygdala (AMY), and 1035 neurons in the medial portion of the mediodorsal thalamus (MDt) using a semi-chronic recording procedure (Figures 1D-F). Neuronal activity differed across areas in different task epochs (Figures S1F-G).

To examine how choices and rewards were represented in each area, we fit the responses of single neurons with a sliding window ANOVA model. The model included multiple task-relevant factors, which were the number of tokens (#token), the change of token numbers (Δtoken), the number of juice drops delivered on cashout trials (#juice), the image pair presented (condition), the stimulus identity (cStim), *a priori* value (cValue, i.e., +2, +1, -1, -2), and direction (cDir) of chosen images. The factors #token, Δtoken, and #juice (Figures 2A-C) were the reinforcement signals that drove the monkeys’ behavior. Neurons in all areas (> 10%), but more in the OFC (> 20%), showed a strong representation of #token (Figures 2A, H). This is consistent with the tokens that were always present on the screen during the trial. There was also a phasic increase in all areas when tokens were updated. The Δtoken was also encoded by neurons in all areas but more strongly in the OFC and VS (Figure 2B). OFC also led the encoding of Δtoken in time (Figure 2B, inset). The #juice activated more than 40% of neurons in every area (Figures 2C, I). OFC played a significant role in encoding the reinforcement signals, having the highest proportion of neurons encoding these task variables. A substantial proportion of neurons in these areas also showed responses to choices, including the identity (Figure 2D), *a priori* value (Figures 2E, G), and direction (Figure 2F) of chosen images. Learning-related values derived from the RW model were encoded consistently across areas (Figure S4B). The stimulus identities, chosen values, and directions were most strongly represented in the AMY, OFC, and MDt, but differences were often subtle. This result shows that task variables related to the choices and rewards were encoded across the ventral network, with different regions encoding specific types of information at different phases.

**Figure 2.**
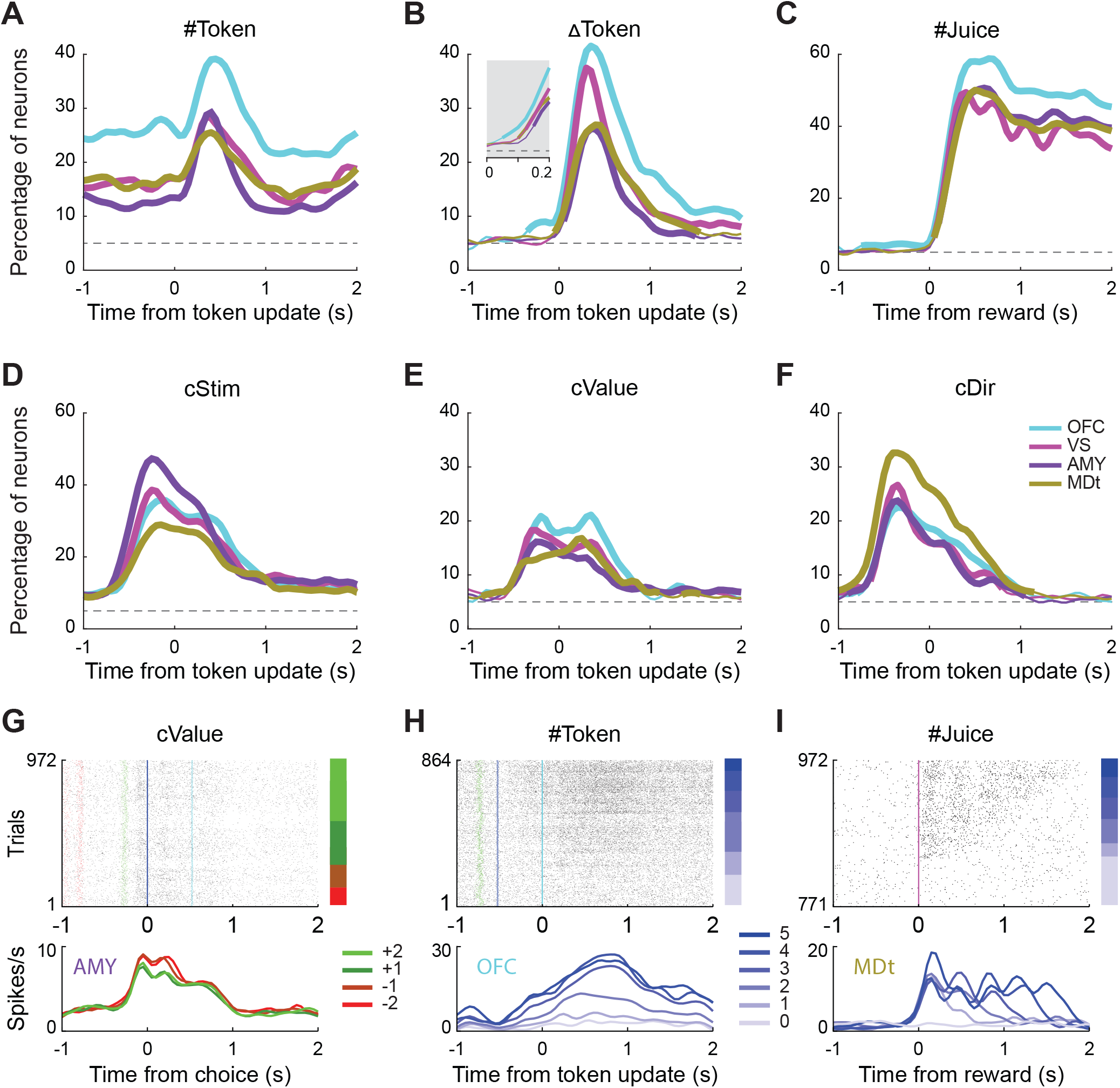
Neural encoding of choices, primary and symbolic reinforcers. (A-F) Percentage of neurons in each area encoding #token (A), Δtoken (B), #juice (C), the identity (D), *a priori* value (E), and direction (F) of chosen images. Data were aligned to either the token update or the onset of juice delivery. Inset in B: ANOVA with 25 ms bin showing response latencies to Δtoken different among areas. Dashed horizontal lines represent the chance level. Thick lines indicate a significant difference between the corresponding area and chance level (binomial test, p < 0.01). (G-I), Example neurons encoded the *a priori* value of the chosen images (G), #token (H), and #juice (I). Each row in the raster plot represents the spikes during a trial. Red, green, blue, cyan, and magenta lines represent fixation, cue onset, choice, token update, and juice delivery. The bars on the right side represent the trial groups under different conditions. The bottom curves indicate mean activities, split by trial groups defined by the bars.

### Diverse encoding of value update information

The updating of value information was represented as gaining or losing tokens. In the first analysis, we used only linear encoding (Figure 2B). However, neurons may encode value information in more diverse ways. To quantitatively characterize the encoding patterns across the areas, we examined the activity of each neuron using a series of linear regression models (Figure S2A). We identified the best-fitting model for each neuron and classified them into different functional categories according to how they were tuned to outcomes.

Here, we classified neurons encoding Δtoken into five types: neurons encoding (1) value linearly across both Gain and Loss (Figures 3A, F), (2) value salience (Figures 3B, G), (3) categorial Gain/Loss (Figures 3C, H), (4) value only for Gain (Figures 3D, I), and (5) value only for Loss (Figures 3E, J). Note that each encoding type could be positively or negatively related to the neural activity. The first category encoded Δtoken linearly (Figure 3A). These neurons encoded the gains and losses on a linear value axis, in other words, processing the gaining and losing of tokens in the same internal value system. Many OFC neurons, but few in the AMY, were of this type. However, more AMY neurons encoded the salience of Δtoken. These neurons represent both gains and losses but with inverse correlations of neural activity and value (Figure 3B). The categorical Gain/Loss signal groups each option as gain or loss, regardless of the magnitude. These neurons were found throughout the ventral network, especially in the OFC and VS (Figure 3C). Many more neurons in these areas encoded gains rather than losses. VS and OFC neurons showed robust and phasic responses to gains (Figure 3D). Only a small proportion of neurons encoded losses, mostly in the OFC (Figure 3E). Before the token update, the chosen images predicted the value update. We also classified the neurons encoding cValue into the same five types (Figure S3A). They showed similar encoding patterns, with activity locked to the onset of the images. This analysis shows that these areas played unique roles in encoding the chosen value and token outcome information.

**Figure 3.**
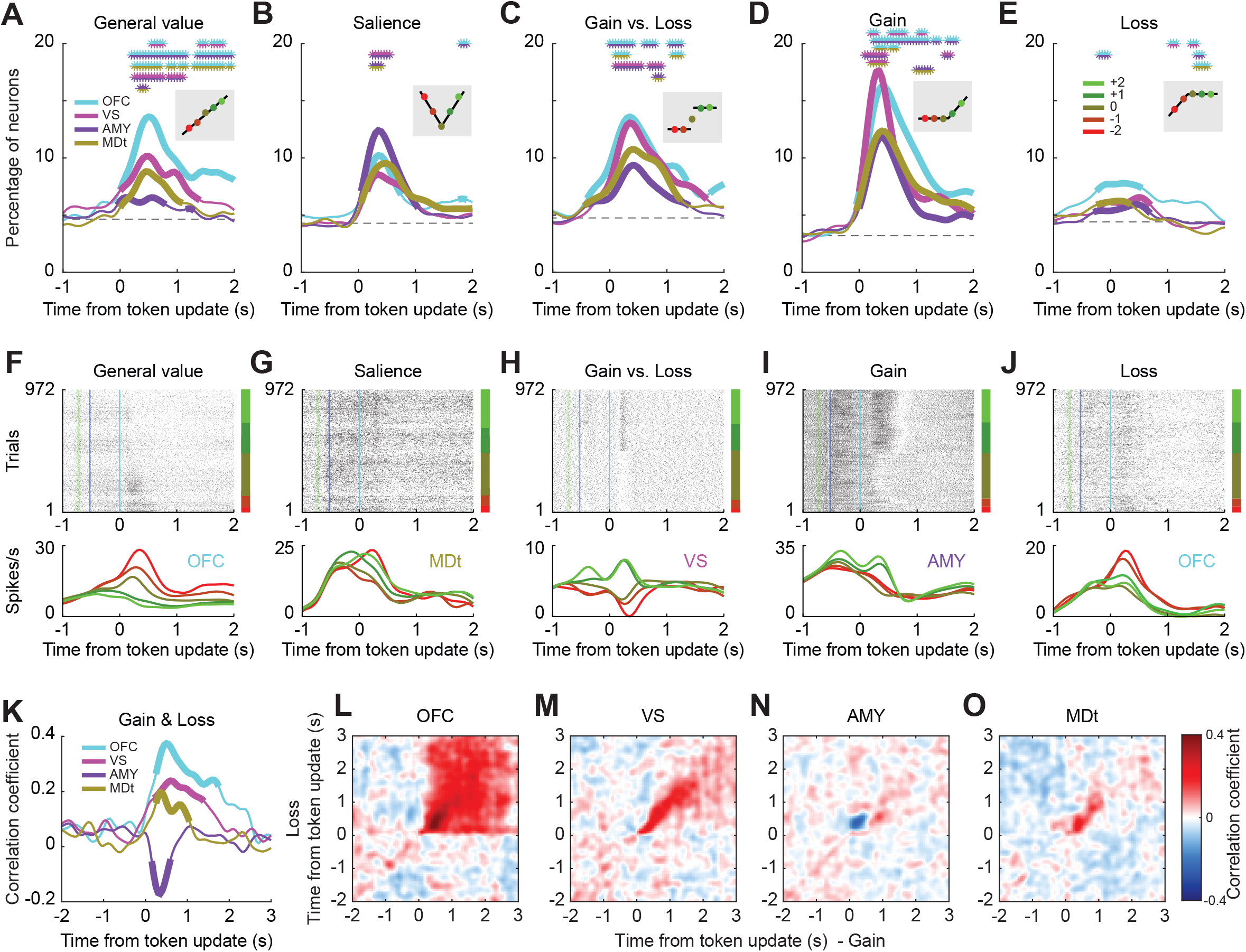
Diverse representation of gains and losses. (A-E) Percentage of neurons in each area encoding Δtoken. The insets indicate the corresponding tuning curves. The coefficients could be positive or negative, meaning the tuning curves could be flipped vertically. (A) General value: monotonic encoding values across gain and loss contexts. (B) Salience: monotonic encoding values across gain and loss contexts but with opposite directions. (C) Gain/Loss: categorical encoding values in gain and loss contexts. (D) Gain: encoding values only in the gain context. (E) Loss: encoding values only in the loss context. Dashed horizontal lines represent chance levels, defined by the mean of all areas during the fixation epoch. Thick lines indicate a significant difference between the corresponding area and chance level (binomial test, p < 0.05). The double-colored asterisks indicate a significant difference between pairs of areas indicated by the colors (chi-square test, p < 0.05). (F-J), Example neurons for each category. (K-O) Co-encoding of gains and losses. Correlations of the Gain and Loss regression coefficients at the same (K) or cross (L-O) time points for each area. Thick lines in (K) indicate a significant difference between the corresponding area and baseline level during the fixation period (binomial test, p < 0.05).

An alternative way to examine the encoding of value update information is to measure the representations of gains and losses separately by running two regressions, either using Gain trials (i.e., Δ*token ∈* [0, 1, 2]) or Loss trials (i.e., Δ*token ∈* [0, −1, −2]), then calculate the correlation of the Gain and Loss regression coefficients (Figures 3K-O). The OFC, VS, and MDt populations showed high co-encoding of gains and losses, which indicates the encoding of gains and losses on similar value axes (Figure 3K). Especially for OFC, the effect was consistent for two seconds after the token update (Figure 4L). This was consistent with the single-cell result (Figure 3A), indicating that OFC processes the gaining and losing of tokens closer to objective outcomes than the other areas (Figure 4K; Fisher’s z-transformation, p < 0.05). Conversely, AMY showed negative correlations between Gain and Loss regression coefficients (Figures 4K, N), indicating population encoding of value salience (Figure 3B).

**Figure 4.**
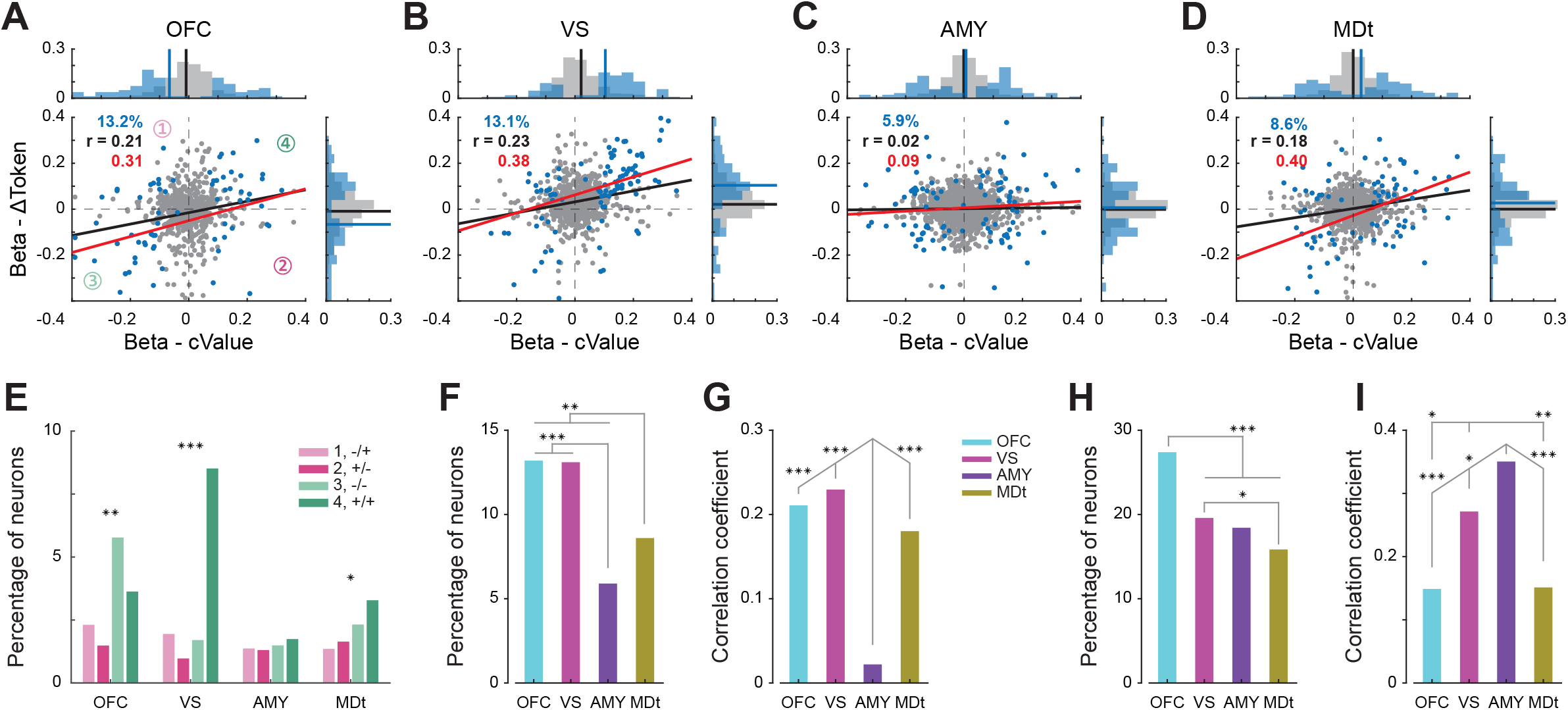
Population encoding of value update and reinforcers. (A-G) Co-encoding of cValue and Δtoken. (A-D) The x- and y-axes represent the regression coefficients of cValue and Δtoken. The blue dots represent the neurons encoding both of them. The gray dots represent the other neurons. Percentages on the left corner indicate the proportions of neurons encoding the cValue and Δtoken. Black and red lines are the lines of best fit for all the neurons and significant neurons, and *r* indicates the corresponding correlation coefficient. The bars summarize the distributions of blue and gray dots, with blue and black lines representing the means. (E) Summary of the distributions of blue dots in each quadrant (also indicated in A) for each area. Chi-square test, *p < 0.05, **p < 0.01, ***p < 0.001. (F) The proportion of neurons encoding cValue and Δtoken in each area. Summary of the percentages in (A-D). Chi-square test, **p < 0.01, ***p < 0.001. (G) Encoding similarity between cValue and Δtoken in each area. Summary of the black *r* in (A-D). Fisher’s z-transformation, ***p < 0.001. Non-significant pairs are not indicated. (H-I) Co-encoding of #token and #juice. (H) The proportion of neurons encoding #token and #juice in each area. Chi-square test, *p < 0.05, ***p < 0.001. (I) Encoding similarity between #token and #juice in each area. Fisher’s z-transformation, *p < 0.05, **p < 0.01, ***p < 0.001.

### Co-encoding of the value update information

In each trial, the monkeys may gain or lose tokens. The value update was predicted by the images (cValue) before the token outcome was delivered (Δtoken). Did neurons in these areas encode the two forms of value information in a similar way? To address this question, we first compared the number of neurons that encoded both forms of value linearly. Neurons encoding cValue in a window 500 ms before the token update and Δtoken in a window 500 ms after the token update were classified as encoding both cValue and Δtoken (blue dots in Figures 4A-D). Fewer neurons in the AMY than in the other areas linearly encoded both factors (Figures 4A-D, F; chi-square test, p < 0.01). To address how similar AMY neurons encoded cValue and Δtoken, we then calculated the correlation of the cValue and Δtoken regression coefficients in each area (Figures S2B-C). AMY neurons showed a significantly lower correlation between cValue and Δtoken than the other areas (Figures 4A-D, G; Fisher’s z-transformation, p < 0.05 for significant neurons, p < 0.001 for all neurons). This result shows that neurons in the AMY encoded the value carried by the images and token outcome less consistently, which is also consistent with AMY primarily encoding salience.

We also found asymmetric encoding of gains and losses in the VS, OFC and MDt (Figure 4E; chi-square test; VS, *x*^2^= 95.74, p < 0.001; OFC, *x*^2^= 19.96, p < 0.001; MDt, *x*^2^= 10.87, p < 0.05). More VS neurons had positive cValue and Δtoken regression coefficients. Together with the results in Figure 3, this indicates that more neurons in the VS fired more when choosing higher-valued images and getting more tokens in the Gain conditions (Figures 4B, E, quadrant 4). OFC neurons showed more balanced encoding in increased and decreased activities (Figure 4E, quadrant 3 vs. 4; chi-square test, *x*^2^= 3.11, p = 0.078). Thus, neurons in the VS tended to respond with increased firing rates to the choice of good options and outcomes, whereas neurons in the OFC had both positive and negative tuning to the same variables.

### Co-encoding of primary and symbolic reinforcers

Although a growing number of studies have adopted symbolic reinforcers, whether they are encoded in the same way as primary reinforcers remains an open question. To address this, we compared the encoding of #token and #juice. We first calculated the number of neurons encoding each variable. Neurons encoding #token in a window 500 ms after the token update and encoding #juice in a window 500 ms after juice delivery were classified as encoding both #token and #juice (Figure S3B). More neurons in the OFC than in the other areas encoded both #token and #juice (Figure 4H; chi-square test, p < 0.001). We then calculated the correlation of the #token and #juice regression coefficients in each area. Surprisingly, neuronal populations in the AMY and VS showed a higher similarity in encoding #token and #juice (Figure 4I and Figure S3C; Fisher’s z-transformation, p < 0.05). This result shows that the symbolic and primary reinforcers were more similarly encoded in the AMY and VS. Although OFC encoded both at the highest level, it did so with different population responses and, therefore, can discriminate these reinforcers best among the areas.

### Value updates in the ventral striatum and orbitofrontal cortex

We used a stochastic reward schedule such that tokens were only updated in 75% of the trials, and token losses only occurred when tokens could be lost. The encoding of Δtoken (Figure 3), therefore, may also include the encoding of RPE (Figure 5A). To address this, we further fit the behaviors with a reinforcement learning model (Figure 1G) and classified each neuron into Δtoken or RPE categories, depending on which variable best described the neuron’s responses, using linear regression models (Figures S4A-B). RPE was calculated with the RW model. Neurons encoding Δtoken responded to the token outcome independently of the learning, whereas neurons encoding RPE encoded the difference between the token outcome and the predicted token outcome, with the prediction estimated by the RW model. Overall, more neurons in the OFC and VS encoded Δtoken or RPE than in the other two areas (Figures S4C-F; chi-square test, p < 0.001). We then split neurons based on whether their regression coefficients were positive or negative (e.g., whether they increased or decreased their firing rate for RPE; Figures 5B-E). For example, a neuron was called a +RPE neuron when classified in the RPE category and with a positive regression coefficient.

**Figure 5.**
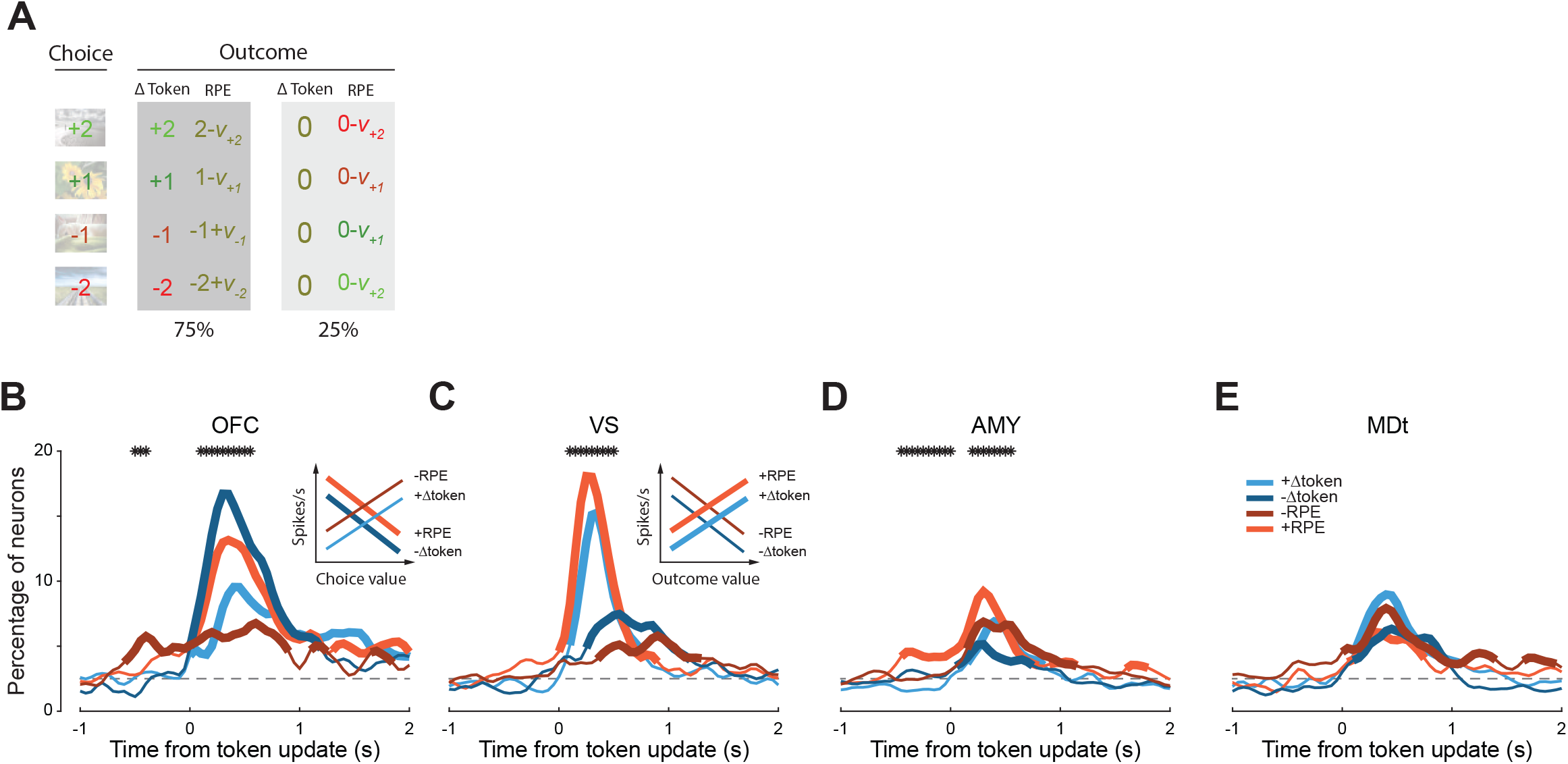
Value updates in the ventral striatum and orbitofrontal cortex. (A) Relationship of *a priori* value, Δtoken, and RPE. *v*_*h*_ indicates the RW model-estimated value of the image with *a priori* value *h*, which got closer to *h* as learning progressed. (B-E) Percentage of neurons in each area encoding +Δtoken, -Δtoken, +RPE, and -RPE. Dashed horizontal lines represent chance levels. Thick lines indicate a significant difference between the corresponding area and chance level (binomial test, p < 0.025). The black asterisks indicate a significant difference among the four categories (chi-square test, p < 0.01). Insets: OFC neurons encoded Δtoken and RPE in a manner relevant to the value of choices. VS neurons encoded Δtoken and RPE correlate with the value of outcomes. Thick lines represent more neurons.

Although a similar proportion of neurons in the OFC and VS encoded token outcome information, they showed different patterns. Most neurons in the OFC at the time of token outcome encoded -Δtoken and +RPE (Figure 5B). In other words, they fired more when losing more tokens (-Δtoken) or when they did not lose tokens after choosing a loss option (+RPE). Specifically, this was aligned with the value gradient carried by the choices (Figure 5B inset), even though it was a signal related to the outcome. On the other hand, most neurons in the VS encoded +Δtoken and +RPE (Figure 5C). Neurons in the VS, therefore, fired more for all good outcomes when gaining or not losing more tokens. This was well aligned with the value gradient carried by the outcomes but not the choices (Figure 5C inset). This suggests that value updates were referenced to outcomes in the VS and to choices in the OFC.

### Population representation of gains and losses

Next, we examined how neural populations in each area dynamically encoded gains and losses at the time of choice and token outcome by measuring population response trajectories^28^. This analysis used pseudo-populations composed of 500 neurons recorded across sessions. We focused on responses in a specific low-dimensional subspace that captured variance due to the eight combinations of cValue and Δtoken (Δtoken equaled cValue or 0). We first defined the axes of the task-related subspace using linear regression coefficients. Then, the condition-averaged population response was projected on each axis to estimate the representation of the corresponding task variables across time (Figure S2D).

The one-dimensional trajectories reflected the dominant tuning at the single-cell level, and correlations between coding of cValue and Δtoken at the population level (Figures 3-5). Trajectories diverged into groups aligned with the value gradient following cue onset for cValue (Figures 6A-D) and following token update for Δtoken (Figures 6E-H). Because all neurons were z-transformed before these analyses, the axes reflect the strength of encoding the corresponding variable. OFC showed the strongest and most balanced representation of positive and negative choices and outcomes among the four areas, with larger divergence among trajectories in both the cValue (Figure 6A) and Δtoken (Figure 6E) axes. The largest deviations for OFC, however, were for Loss conditions. OFC trajectories also showed the population representation of RPEs, with zero token outcomes for Loss options intermediate between Loss and Gain outcomes and zero token outcomes for Gain options merged with Loss outcomes (Figure 6E). Because cValue and Δtoken were correlated in OFC, the cValue trajectories also crossed following the token update (Figure 6A).

**Figure 6.**
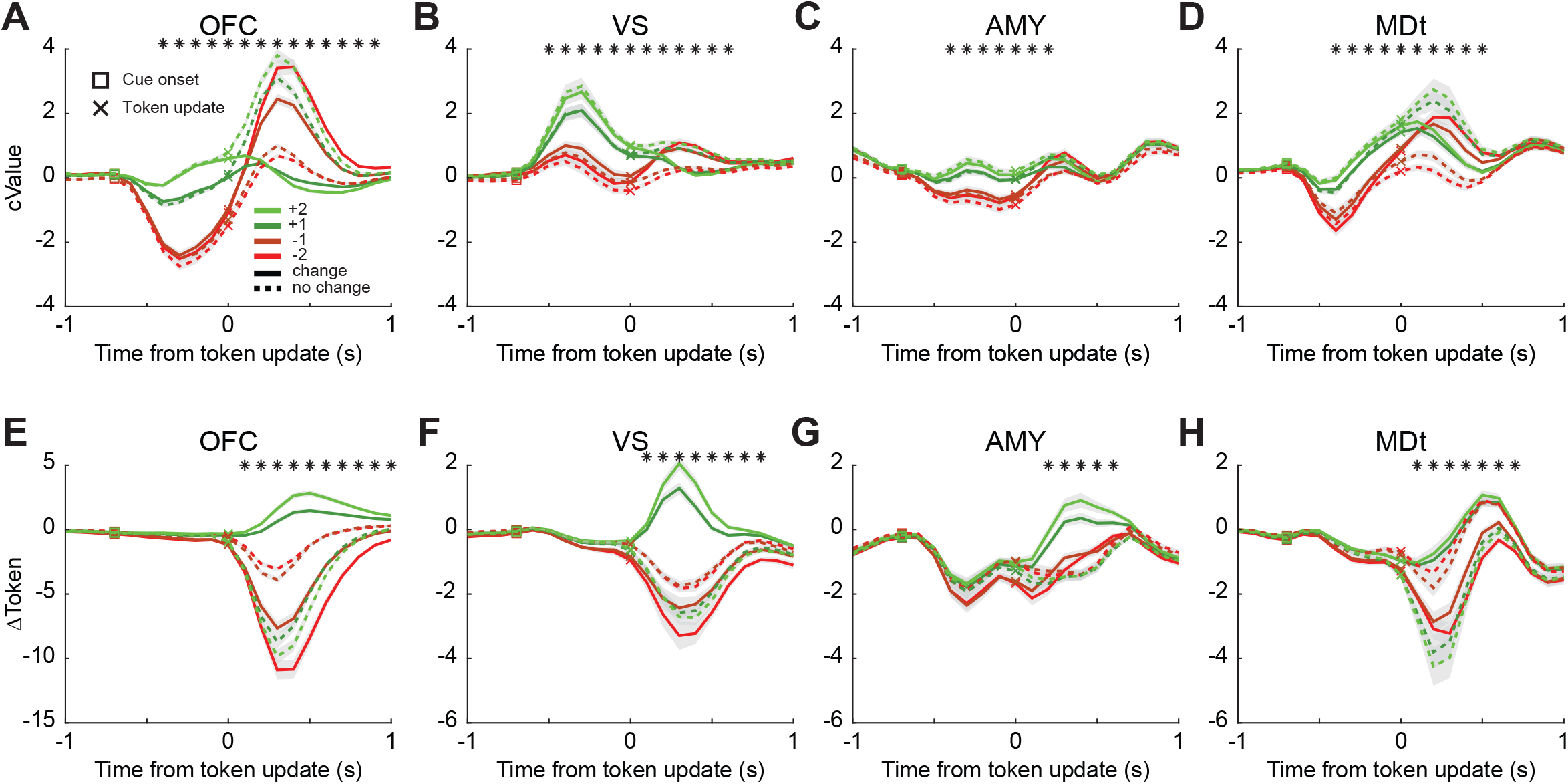
Population representation of gains and losses. (A-D) Projections of neural population activity on the cValue axis in the task-variable-specific one-dimensional subspace. The y-axis indicates the value of the projection, divided by the number of neurons (always 500). All neural activity was also z-scored. Therefore, the y-axis represents the average z-scored population deviation per cell. Trials were grouped by cValue and outcome. Solid lines indicate the conditions when tokens were delivered, and dashed lines indicate the conditions in which tokens were not delivered. The analysis was repeated 1000 times by randomly choosing 500 neurons from each population without replacement. Shaded zones show the mean ± SD across iterations. The squares and crosses represent the cue onset and token update. The black asterisks indicate a significant difference among the eight conditions (bootstrap test, p < 0.01). (E-H) Projections of neural population activity on the Δtoken axis in the task-variable-specific one-dimensional subspace.

VS population activity also discriminated gains and losses well but with a bias to the Gain conditions. The Gain trajectories had steeper peaks (Figure 6B) and were better separated (Figures 6B, F) than the Loss trajectories. The VS also showed the crossover in cValue trajectories after outcomes (Figure 6B). AMY trajectories were grouped into Gain and Loss groups for cValue (Figure 6C), indicating the overall population coding of Gain vs. Loss. However, the Δtoken trajectories, about 250 ms after the token update (Figure 6G), had the weakest responses for all zero token outcomes, and stronger responses for both gains and losses, indicating the overall population coding of the salience of Δtoken. MDt trajectories showed similar patterns to those of OFC, including the crossover following zero token outcomes. This suggests that they shared similar information. Overall, these results are consistent with the results shown in Figures 3-5, which confirm the unique contributions of each area in encoding gains and losses.

### Flow of token information in the ventral network

Brain areas function as part of big networks but not as individual isolated areas. The task-relevant information also is not isolated in each area. To understand the flow of token information within the network, we measured the linear dynamics of the trajectories within a trial. This analysis used only simultaneously recorded ensembles and was carried out trial-by-trial. First, we projected the population neural activity from single trials into a 3-D subspace to generate the population state-space representation of #token, Δtoken, and cDir (Figure S2D). Then, we put these task-variable-specific latent variables together into a matrix *X*. The rows included all the latent variables from all the areas, and the columns included each trial (Figure 7A). We stacked all the sessions (N = 16) with more than ten neurons recorded from each area simultaneously, then estimated the loading matrix *A* (Figure 7B). The matrix *A* characterized the flow of information among variables and areas (Figure S2E), with its columns representing the source areas and variables, and the rows representing the target areas and variables (Figure 7B). The matrices were estimated separately at each point in time, as a local linear approximation to the dynamics, which were likely nonlinear.

**Figure 7.**
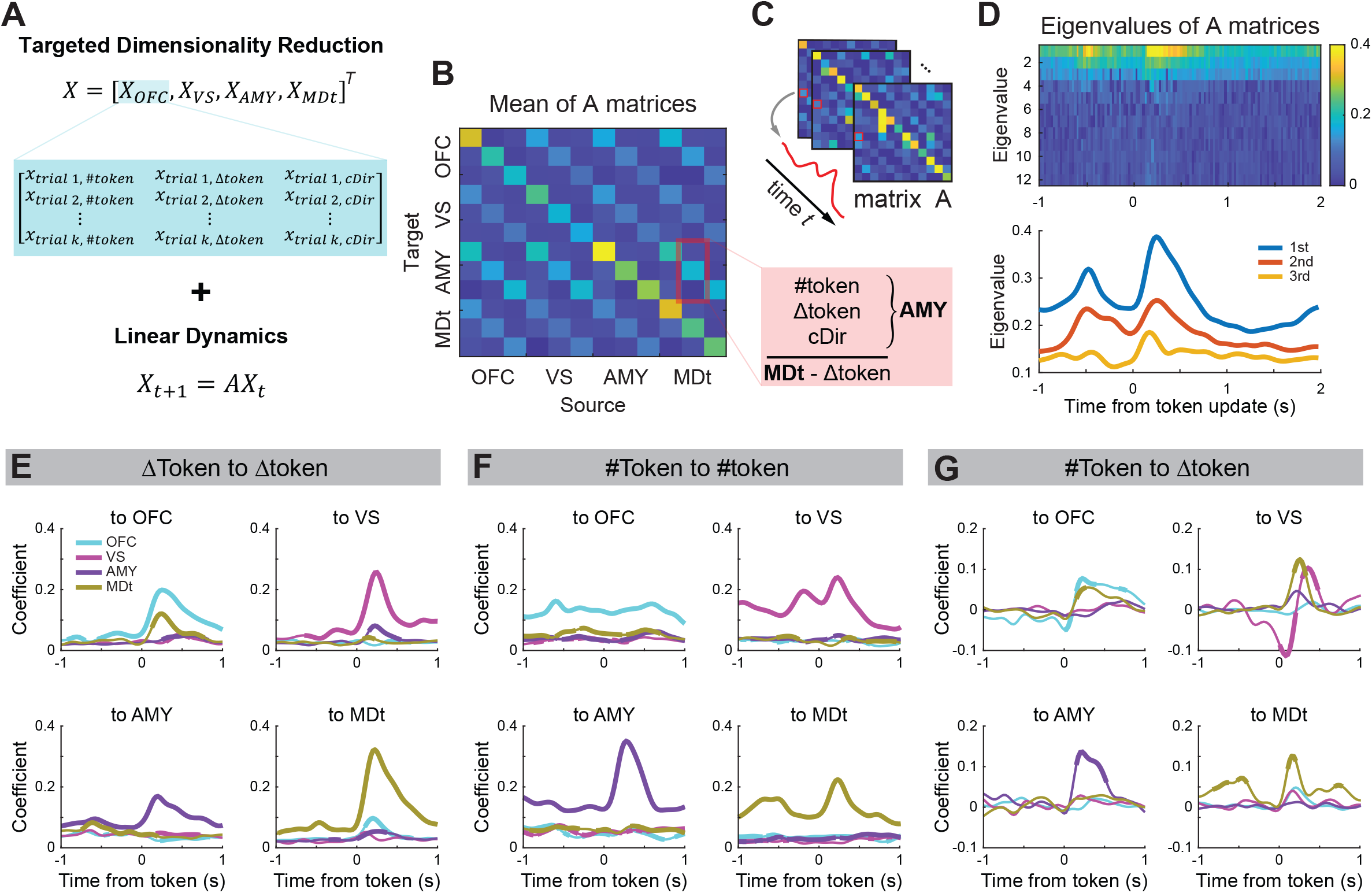
Flow of token information in the ventral network. (A-D) Detecting network communications with low dimensional information dynamics. (A) The task-related population response matrix consisted of single-trial-level population projections. Linear dynamic regression measured time-dependent flows of information across task variables and brain areas. (B) The loading matrix *A* maps the relationship between rows in matrix *X*. The x-axis represents the source areas and variables, and the y-axis represents the target areas and variables. For example, the values indicated by the red square represent the flow of Δtoken information from the MDt to #token, Δtoken, and cDir information in the AMY. (C) The continuous values in *A* matrices represent the strength of information flow across task variables and brain areas along the time course of the trial. (D) The eigenvalues of *A* matrices capture the strength of the overall information flow among the areas in the network. (E-G) The flow of token information across areas, including within the Δtoken (E), #token (F), and between them (G). The titles indicate the source and target task variables. Each panel indicates one target area, and lines indicate source areas. Thick lines indicate a significant difference between the corresponding area and shuffled data (two sides z-test, p < 0.01). Shuffling was done by keeping the same conditions but shuffling the trial order of matrix *X*.

The eigenvalues of the *A* matrices captured the time constant of the dynamics of information flow among the areas in the network. The top two eigenvalues, which show two peaks, indicate robust information flow during the cue and token update epochs, and the third eigenvalue captured information flow during the token update epoch (Figure 7D). The left and right eigenvectors of the matrix *A* capture the temporal dynamics of output (Figures S5A-C) and input (Figures S5D-F) information. The first eigenvector also reflected the temporal structure of the task, showing phasic activity that was locked to the associated task variables. The values in the *A* matrices indicate the strength of information flow between a specific pair of areas and task variables. The temporal dynamics of the information flow varied across the time course of the trial (Figure 7C). We examined the flow of token information across areas and task variables (Figure S6) and found the strongest information flow existed within the same area for most conditions. This is consistent with other work showing that within-region dynamics are higher dimensional than across-region dynamics ^29^.

Our dynamics analysis showed good specificity between task variables. For example, despite the fact that cDir, #token, and Δtoken were represented across areas, there were minimal interactions between cDir and the value signals (Figures S6F-H). Aiming to address the flow of token information, we focused on the interactions within and between the Δtoken and #token. For Δtoken (Figure 7E), information flows showed peaks phase-locked to the token update. There was strong reciprocal information flow between the OFC and MDt, reflecting the underlying anatomy^19^. We also found flow of information from AMY to the other areas, especially VS. For the #token (Figure 7F), the information flow was relatively continuous along the time course of the trial, which is consistent with the tokens being present on the screen across trials.

More interestingly, there was a temporal derivative in the flow of information from #token to Δtoken in the OFC and VS (Figure 7G, 100 ms before vs. after derivative; Fisher’s z-transformation; VS to VS, p < 0.001; OFC to OFC, p < 0.05). This shows the mechanism by which Δtoken was calculated in the network. Specifically, OFC and VS computed the difference in #token following and prior to the choice (i.e.,

Δ*token* = *#token*_*after*_ − *#token*_*after*_). Also, we found a stronger flow of information from the #token to Δtoken than from the Δtoken to #token (Figures S6B, D; Fisher’s z-transformation, p < 0.05). This suggests that the updating of values in this four-area network originated predominantly in the VS and OFC, consistent with the single-cell results (Figures 5B-C). Because both areas showed time-derivative dynamics, it seems likely that they may both be calculating the update. However, the derivative in the OFC appeared about 100 ms earlier than in the VS, similar to the time of single neurons in the OFC leading Δtoken encoding relative to the VS (Figure 2B inset). During this calculation, phasic reciprocal information flow also existed between OFC-MDt and MDt-VS (Figure 7G), indicating that MDt bridged the information flow from the OFC to VS.

## DISCUSSION

We examined the single-cell, population, and network representation of token-based RL across the ventral cortico-striatal-thalamo-cortical network^1,19,30^. We found that the OFC played an important role in computing information about token updates. While all areas coded this information, OFC coded it earlier and more strongly. We also found that the other areas contributed unique computations. The VS showed a strong bias towards encoding appetitive choices and outcomes, including positive RPEs, with increased firing rates. On the other hand, OFC showed a more balanced encoding of positive and negative choices and outcomes with increased and decreased firing rates. OFC also strongly encoded primary and symbolic reinforcers but distinguished them at the population level. AMY, on the other hand, encoded both reinforcers similarly at the population level. AMY also showed enhanced salience coding and did not show correlations between choice and outcome value coding. Finally, MDt showed enhanced coding of actions and mediated information flow between the OFC and VS during token update calculations.

### Primary and symbolic reinforcers

Human studies typically use symbolic reinforcers, while animal studies use primary reinforcers. However, similarities and differences between the population coding of primary and symbolic reinforcers have not been examined. AMY and, to some extent, VS populations showed stronger correlations between primary and symbolic reinforcers and, therefore, encoded tokens and primary reinforcers similarly. This may explain why earlier lesion work found AMY linked conditioned stimuli to the specific reward properties of the unconditioned stimuli they predicted^31^. OFC showed the most substantial coding of both primary and symbolic reinforcers but encoded them with the lowest similarity, therefore discriminating them well. Thus, OFC represents value information with high fidelity, using a code that preserves detailed information about the type and valence of reinforcement^32^.

### Gains and losses

We found that VS showed a strong bias towards monotonically encoding gains, specifically coding rewarding choices and outcomes, and better than the expected outcomes, with increased firing rates. Thus, VS consistently signaled token gains with increased firing rates. This is consistent with human imaging studies^33^ and the longstanding suggestion that VS is important for motivation^34,35^. AMY neurons showed enhanced tuning for salience. The AMY population, uniquely among the areas, did not show a correlation between linear choice value and outcome value encoding, and showed negative correlations, at the population level, between gain and loss encoding, both of which are also consistent with salience coding.

Previous studies found AMY responses to appetitive and aversive stimuli but could not assess salience because they only used single outcomes for each valence^3,36,37^. OFC showed monotonic encoding of value across both positive and negative outcomes. Unlike VS and AMY, OFC neurons encoded values with both positive and negative slopes with a slight bias towards neurons responding more for negative value choices and loss outcomes. Recordings in the pregenual anterior cingulate cortex (ACC)^38^ and the insula^8^ similarly found neuronal populations coding appetitive and aversive choices with both positive and negative slopes. Thus, cortical areas show a more balanced coding of gains and losses than the VS and AMY.

### OFC contributes to the encoding of symbolic reinforcers

We found that OFC encoded accumulated tokens and changes in tokens at both the single-cell and population levels. These findings are consistent with previous work showing that OFC codes state information about the environment^39-42^, as tokens and token updates define states and state transitions in this task^43^. State representation is also referred to as a cognitive map, particularly when states have to be inferred^44,45^. It is a map because states are nodes on graphs, and one has to know the current state and subsequent states to which one can transition. Within reinforcement learning, states are the environmental variables relevant to the learning and action selection process^46^. We also found that OFC encoded choice value, which is consistent with previous work showing that OFC codes economic value^47^. However, we have found that choice value was broadly encoded across our network, and OFC did not encode value more strongly than other areas. We did find differences in the way in which OFC encoded values relative to the other areas, as discussed above. OFC has a high-fidelity representation of the state that includes negative outcomes and the strongest encoding of accumulated tokens.

### The calculation of value updates

We also examined network computations within targeted information dimensions. Analysis of single-trial communication using neurons recorded from multiple regions simultaneously can provide evidence of dynamic network processes that underlie behaviors^48^. Most studies that have examined interactions between areas with neurophysiology data have focused on pairwise interactions between cortical regions and have also not identified dynamic signatures of computations. Instead, they have shown that interactions occur within specific dimensions and often during particular periods within a task. For example, an early study found that choice-relevant signals were relayed from the prefrontal to the parietal cortex to guide behavior^49^. A more recent study found that stronger value coding in the OFC led to accelerated ramping of signals in the ACC^50^. We recently identified object-to-direction information flow among lateral prefrontal cortex subregions in a task in which the location of a valuable stimulus had to be identified before a saccade could be directed toward it to make a choice^51^.

The current study measured the flow of value information using simultaneous population recordings from four areas in the ventral network. This allowed us to control for multiple, although not all, inputs to each area. We found that token update information was calculated in the OFC and VS as a time-derivative of the information about accumulated tokens (i.e., Δ*token*_*t*+1_ = *#token*_*t*+1_− *#token*_*t*_). The signal was earlier in the OFC but stronger in the VS. Thus, the change in accumulated tokens drove information about token outcomes. This is consistent with the finding that OFC and VS strongly encoded Δtoken and token prediction errors at the single-cell level. Note that in previous work with probabilistic reward outcomes, we did not find strong encoding of RPE in these structures^16,44^. We found minimal support for the alternative hypothesis that token outcomes were integrated to generate information about accumulated tokens (i.e., *#token*_*t*+1_ = *#token*_*t*_ + *Δtoken*_*t*_). Thus, analysis of neural dynamics identified an explicit computational correlate of a behavioral process. We also found that MDt mediated interactions between the OFC and VS with respect to token updates by mediating reciprocal information flow between OFC-MDt and MDt-VS. Interestingly, we did not identify a direct interaction between the OFC and the VS, even though they are connected directly^52,53^. This suggests that the token updates were mediated within the cortical-thalamic-striatal circuitry but not the corticostriatal circuitry.

### Effects of loss of specific nodes on reinforcement learning

These results also provide insight into the task-dependent effects of lesions in previous work. For example, tasks that use probabilistic delivery of primary reinforcers have shown deficits following lesions to all nodes of the ventral network^14,15,17,18,54,55^. However, lesions of the AMY have almost no effect on learning in the tokens task, and lesions of the VS only affect learning to discriminate gain magnitude but not gains vs. losses ^7,23^. Here, we found that AMY showed enhanced salience coding. Salience coding can be used as a learning signal in the context of probabilistic reward outcomes because salience and gains relative to no outcome are equivalent. However, salience coding cannot be used for learning in token-based reinforcement because large gains and large losses are represented similarly. This may also explain why we found no evidence for the effects of AMY lesions on unsigned prediction error variables in Pearce-Hall models of RL that had been reported previously^56^ when examined in probabilistic reward tasks^17^. We also found that VS was strongly biased toward encoding appetitive outcomes. Therefore, lesions to the VS would be expected to have a larger effect on discriminating among gains, as opposed to between gains and losses^7^. Although we did not record in the insula, recent work has shown that the insula has substantial coding of losses in token-based decision-making^8^, consistent with early fMRI results^57^. Therefore, it is possible that manipulations of the insula would show effects specific to loss outcomes. Finally, we found that OFC may preferentially mediate the calculation of token reinforcement, and therefore, manipulations of OFC may have large effects on learning in the tokens task.

## Conclusion

Information about task variables was represented across the ventral network. Although all areas represented task variables, they did so differently. AMY encoded outcome salience and encoded the primary and symbolic reinforcers similarly. VS and OFC encoded value information, with VS strongly biased towards coding positive outcomes with increased firing rates and OFC more balanced towards coding positive and negative outcomes. OFC and VS calculated Δtoken as the time-derivative of accumulated tokens, with a shorter latency in the OFC. MDt mediated the interactions about token updates between the OFC and VS during this process. Importantly, the representation and computation of symbolic reinforcement appear to be more strongly mediated by cortical structures, in this case, OFC, than subcortical structures, which may differ from primary reinforcement.

## ACKNOWLEDGMENTS

We thank Christos Constantinidis, Vincent Costa and Diana Burk for their valuable comments, as well as Andy Mitz, Craig Taswell, Miriam Janssen and Sarah Falkovic for their technical help. This work was supported by the Intramural Research Program of the National Institute of Mental Health (ZIA MH002928) to B. B. A. and BBRF NARSAD Young Investigator Award (30892) to H. T. Anatomical MRI scanning was carried out in the Neurophysiology Imaging Facility Core (NIMH, NINDS, NEI).

## AUTHOR CONTRIBUTIONS

Conceptualization, R. B., H. T., and B. B. A.; Methodology, H. T., R. B., and B. B. A.; Investigation, H. T., and R. B.; Visualization, H. T., and B. B. A.; Writing – Original Draft, H. T., and B. B. A.; Writing – Review & Editing, H. T. and B. B. A.; Funding Acquisition, H. T. and B. B. A.; Supervision, B. B. A.

## STAR METHODS

### KEY RESOURCES TABLE

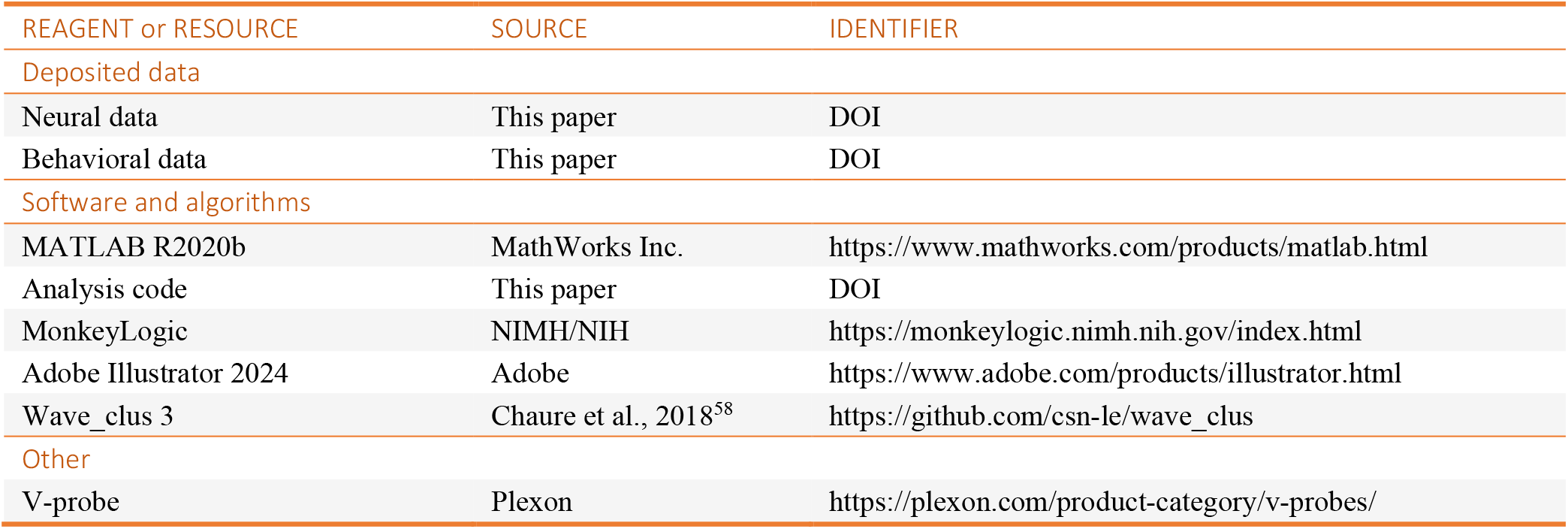

| REAGENT or RESOURCE | SOURCE | IDENTIFIER |
| --- | --- | --- |
| **Deposited data** |  |  |
| Neural data | This paper | DOI |
| Behavioral data | This paper | DOI |
| **Software and algorithms** |  |  |
| MATLAB R2020b | MathWorks Inc. | https://www.mathworks.com/products/matlab.html |
| Analysis code | This paper | DOI |
| MonkeyLogic | NIMH/NIH | https://monkeylogic.nimh.nih.gov/index.html |
| Adobe Illustrator 2024 | Adobe | https://www.adobe.com/products/illustrator.html |
| Wave_clus 3 | Chaure et al., 2018<sup>58</sup> | https://github.com/csn-le/wave_clus |
| **Other** |  |  |
| V-probe | Plexon | https://plexon.com/product-category/v-probes/ |

RESOURCE AVAILABILITY

#### Lead contact

Further information and requests for resources and reagents should be directed to and will be fulfilled by the lead contact, Bruno B. Averbeck.

#### Materials availability

This study did not generate new unique reagents.

#### Data and code availability

The datasets supporting the current study will be publicly available as of the date of publication. DOIs are listed in the key resources table.

All original codes will be publicly available as of the date of publication. DOIs are listed in the key resources table.

Any additional information required to reanalyze the data reported in this paper is available from the lead contact upon request.

### EXPERIMENTAL MODEL AND STUDY PARTICIPANT DETAILS

#### Subjects

The experiments were performed on one adult male (10 kg) and one adult female (7.5 kg) rhesus macaque (*Macca mulatta*). They were 8-10 years old. The monkeys were pair-housed when possible and had access to food 24 hours per day. On testing days, the monkeys were placed on water control and earned their juice through performing the task. On non-testing days, the monkeys were given *ad libitum* access to water.

#### Ethics

Experimental procedures for all monkeys followed *the Guide for the Care and Use of Laboratory Animals* and were approved by the National Institute of Mental Health Animal Care and Use Committee.

### METHOD DETAILS

#### Experimental Setup

Monkeys were trained to perform a saccade-based two-armed bandit task. Stimuli were presented on a 19-inch LCD monitor situated 40 cm from the monkeys’ eyes. During training and testing, the monkeys sat in a primate chair with their heads restrained. Stimulus presentation and behavioral monitoring were controlled by MonkeyLogic^59^. The eye movements were monitored at 400 fps using a Viewpoint eye tracker (Arrington Research, Scottsdale, AZ) and sampled at 1 kHz. A fixed amount of apple juice was delivered through a pressurized plastic tube gated by a solenoid valve on rewarded trials.

### Task Design

The task was developed and first used in our previous study ^7^. Each session had nine blocks. Each block used four novel images associated with different values, including +2, +1, −1, and −2. The monkeys obtained more tokens by choosing images associated with larger values. Choosing one of the images led to gaining or losing a corresponding number of tokens. Also, token numbers could not be negative, so choosing a loss image when there were no accumulated tokens had no effect. We used a stochastic reward schedule. The number of tokens updated 75% of the time and did not change 25% of the time. To complete a trial successfully, the monkey first acquired and held central fixation for 400-600 ms. Then, two of the four images were randomly selected by the computer and displayed on the screen. The animal made its selection by saccading to one of them. The unchosen image disappeared when the monkey reached the target. Saccade fixation was maintained on the chosen image for 500 ms. After that, the chosen image also disappeared, and the token number was updated according to the chosen image. Tokens accumulated across trials and were cashed out for juice every four to six trials, with the interval randomly selected. At cash-out, the animals were given one drop of juice for each token. When each drop of juice was delivered, one token was removed from the screen. Up to 12 tokens were accumulated and were displayed on the screen across trials.

The task had six individual conditions defined by the possible values of the image pairs. The conditions within a block of 108 trials (6 conditions × 2 counterbalanced for left and right sides × 9 repetitions) were presented pseudo-randomly. The animals saw each condition twice, once on the left and once on the right, every 12 trials before seeing any condition a third time. We introduced four novel images at the beginning of each block. Images provided as choice options were normalized for luminance and spatial frequency using the SHINE toolbox for MATLAB, described previously^7^.

#### Surgical procedures

Each monkey was surgically implanted with a titanium headpost, and a 25 × 35 mm recording chamber to allow vertical grid access to the OFC, VS, AMY, and MDt (Figures 1D-F). Grid holes for the MDt had 16° angles to allow better access to the target area (Figure S1D). Chamber placements were planned and verified with T1 and T2 magnetic resonance imaging (MRI, 3.0 T). Small burr holes were drilled above each target area. A grid was installed, and one guide tube was inserted through each burr hole. The guide tubes were lowered to about 1-4 mm (depending on the target areas) above the target areas and were glued to the grid. We removed the guide tubes and placed new ones in the adjacent locations after 3-5 recording days (Figures S1D-E). The locations of the guide tubes were verified with MRI after guide tube replacement. All sterile surgeries and MRI scans were performed under anesthesia.

#### Neurophysiological Recordings

Neurophysiology recordings (Figures S1C-D) began after the monkeys had recovered from the surgery. We lowered one linear electrode array (V-probe, Plexon Inc, Dallas, TX) into each guide tube on every recording day. Thirty-two channel electrodes with 150 μm inter-contact spacing probes were used in the VS and MDt, and 64-channel electrodes with 150 μm inter-contact spacing probes were used in the OFC and AMY. The probes were advanced to their target location by a four-channel micromanipulator (NAN Instruments, Nazareth, Israel) attached to the recording chamber. The depths of the neurons were estimated by their recording locations relative to the tip of the guide tubes (verified with MRI). Electrophysiological data were acquired with a 512-channel Grapevine System (Ripple, Salt Lake City, UT). The spike acquisition threshold was set at a 4.0 × root mean square (RMS) of the baseline signal for each electrode. Behavioral event markers from MonkeyLogic and eye-tracking signals from Viewpoint were sent to the Ripple acquisition system. The extracellular signals were high-pass filtered (1 kHz cutoff) and digitized at 30 kHz to acquire the single-cell activity. Spikes were sorted offline via Wave_clus 3^58^.

#### Choice behavior

Each block had six conditions. The conditions pseudo-randomly appeared in the task. The number of observations in each condition increased as a function of learning. We quantified choice behavior during the task by measuring the fraction of choosing the image associated with a higher value in each condition. We then measured how different task variables affected the choice behavior by fitting a multi-way ANOVA model. Factors, including the number of observations (Trial), Gain/Loss condition (Loss/Loss, Loss/Gain, or Gain/Gain), value difference of the options (Δvalue, e.g., Δvalue = 4 in the condition of -2 vs. +2), the number of tokens before choice (#token), and the number of trials since last token cashout (CashID) were used in the model. The model was run session by session to measure the contribution of each variable in each session, using all the sessions to acquire the statistics for each variable. All trials in which monkeys chose one of the two stimuli were analyzed. Trials in which the monkey broke fixation, failed to make a choice, or attempted to saccade to more than one target were excluded.

#### Effect size

To quantify the contribution of each task variable to the monkey’s choice. We computed each factor’s effect size, *ω*^2^, from the ANOVA model output. *ω*^2^is an unbiased estimator of the amount of variance in neural activity explained by each task variable, and ranges between -1 and 1^60^. It is given by:

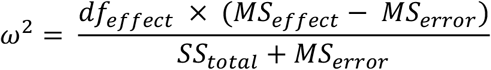

where *df*_*effect*_ refers to the degrees of freedom associated with the factor, *MS*_*effect*_ refers to the mean squares, *MS*_*error*_ is the mean squared error, *S*_*total*_ is the sum of squares of all factors.

#### Responsive neurons

To identify neural responses to different task components, we fit a sliding window multi-way ANOVA model to spike counts computed in 200 ms bins, advanced in 50 ms increments, and time-locked to token update. Factors, including the number of tokens on the screen (#token, may change after the choices), the change of token numbers (Δtoken), the drops of juice delivered (#juice), image pairs with different value combinations presented (condition, e.g., +2 vs. -1), the order of blocks (blockID), the identity (cStim), *a priori* value (cValue), and direction (cDir) of chosen images were used in the model. The stimulus identity was the specific image used in each block to represent each outcome, which was the interaction of cValue and blockID. The factor blockID was used to remove non-stationarity due to drift.

#### Statistic test for the proportion of neurons

The binomial test was applied to test whether the proportion of responsive neurons was significantly above the chance level (5% most of the time). The chi-square test was used to compare the proportions of responsive neurons between different pairs of brain areas or among four areas. Significant encoding for each factor at each time bin was evaluated at p < 0.05. A neuron that showed a significant response to a factor in no less than three contiguous bins in the statistics was considered to be responsive to that factor.

#### Neuronal coding regression analysis

To quantitatively characterize how the updating of values (including cValue and Δtoken) was encoded in different areas, we examined the activity of each neuron using a series of multivariate linear regression models^8^. The dependent variable, neural activity, was first z-scored by subtracting the mean response from the firing rate at each time and in each trial and dividing the result by the standard deviation of the responses. Both the mean and the standard deviation were computed by combining the neurons’ responses across all trials and times. We then described the z-scored responses of neuron *i* at time *t* as a linear combination of several task variables. The independent variables were the same as the factors used in the ANOVA model:

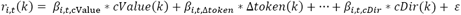

where *r*_*i,t*_ (*k*) is the z-scored response of neuron *i* at time *k* and on trial *t*, Δ*token*(*k*) is the change of tokens on trial *k*. The regression coefficients, *β*_*i,t,f*_, describe how much the trial-by-trial firing rate of neuron *i*, at a given time *t* during the trial, depends on the corresponding task variable *f*.

We tested all potential combinations of tuning for Δtoken (six forms: General value, Salience, Gain/Loss, Gain, Loss, nan) and cValue (six forms: General value, Salience, Gain/Loss, Gain, Loss, nan). Nan means the variable was removed from the model. A particular variable could only be represented by one specific form but not by combinations of more than one form (e.g., a model containing the Δtoken variable could include Gain or Salience but not both). In total, we tested 6 × 6 = 36 models for each neuron. This method is also illustrated in Figure S2A. We determined the best-fitting model for each neuron using the Akaike information criterion (AIC). Neurons were classified into different functional categories (up to one category per variable) according to the combination of the forms of these two variables included in the best-fitting model. We computed the AIC:

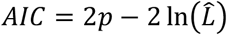

where *p* is the number of free parameters in t hemodel, and 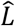 is the maximized value of the likelihood function.

#### Population coding similarity

The regression coefficient, *β*_*i,t,f*_, from the multivariate linear regression model reflects the weight of one task variable, *f*, in explaining the variation of the neuron’s activity at time *t*. The regression coefficients,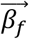, from a neural population, represent the weights of each neuron in the population encoding one task variable. Hence, computing the correlation between two regression coefficient vectors from the same neural population tells us how similar a neural population encodes two task variables (Figures S2B-C):

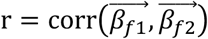

Where 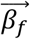 is a vector that consists of regression coefficients of task variable *f* in a neural population. To assess the significance of the correlation between two regression coefficients, we transformed the correlation coefficients into normally distributed z-scores using Fisher’s z-transformation.

#### Reward prediction error

The RPE was defined as the difference between the change of tokens Δ*token*(*t*) and the estimated value of the chosen image, which is given by:

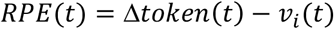

where *t* means trial order, and *i* means the chosen option among two options. The updating of value *v*_*i*_ was estimated using the Rescorla–Wagner equation:

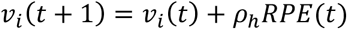

where *ρ* is the learning rate. The updated value estimate *v*_*i*_ (*t* + 1) equals the previous value estimate *v*_*i*_ (*t*) plus the RPE scaled by the cue-dependent learning rate *ρ*_*h*_ for images associated with different *a priori* values *h*. These values were then passed through a soft-max function to give choice probabilities for the image pairs presented in each trial:

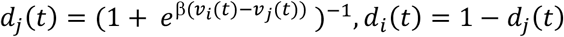

*β* is the choice consistency or inverse temperature parameter, fit across all six cue conditions, and *i* and *j* are the two choice options. We then maximized the likelihood of the animal’s choices, *D*, given the parameters, using the cost function:

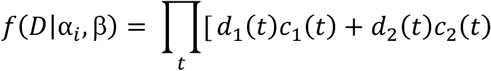

where *d*_1_(*t*) is the choice probability value for one option on trial *t*, and *c*_1_(*t*) and *c*_2_(*t*) are indicator variables that take on a value of 1 if the corresponding option was chosen, and 0 otherwise. This model was fit across blocks in each session for each monkey to give one set of free parameters for each session.

#### Targeted dimensionality reduction

We used the regression coefficients described above to identify dimensions in state space representing each task variable^28^. For each variable, we first build a set of coefficient vectors 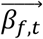 whose entries *β*_*f,t*_ (*i*) correspond to the regression coefficient for task variable *f*, time *t*, and neuron *i*. The vectors 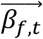 (of length *N*_*unit*_) are obtained by simply rearranging the entries of the vectors 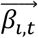 (of length *N*_*f*_) computed above. Each vector,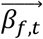, thus corresponds to a direction in state space that accounts for variance in the population response at time *t*, due to variation in task variable *f*.

We used time-dependent regression vectors, *B*_*t*_, which is a matrix with each column corresponding to the one regression vector 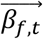. We referred to them as the ‘task-related axes’. These axes span a ‘regression subspace’ representing the task-related information the neural population coded. We then projected the single-trial population responses onto these axes to study the representation of the task-related variables in each trial:

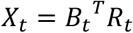

where *R*_*t*_ is single-trial population response at time *t*, which is of dimension *N*_*unit*_ × *N*_*trial*_. And *X*_*t*_ is the set of time-series vectors over all task variables and trials, which is of dimension *N*_*f*_ × *N*_*trial*_. It represents the population coding of task-related information at time *t* of every single trial. This method is also illustrated in Figure S2D.

#### Linear dynamics

We fit an autoregressive, linear model to the *X*_*t*_,

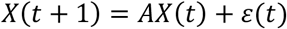

where *X*(*t*) = [*X*_*OFC*_ (*t*), *X*_*VS*_(*t*), *X*_*AMY*_(*t*), *X*_*MDt*_ (*t*)]^*T*^ is a matrix of dimension (*N*_*area*_· *N*_*f*_) × *N*_*trial*_, and *X*_*area*_ (*t*) = [*x*_*k,#token*_(*t*), *x*_*k*,Δ*token*_(*t*), *x*_*k,cDir*_ (*t*)] is a matrix of dimension *N*_*trial*_ × *N*_*f*_. *ε*(*t*) is the noise term. *A* is a 12 × 12 dynamics matrix, where the 12 = 4 × 3 dimensions correspond to the combinations of brain areas and task variables. The model was fit using data points in a non-overlapping 25 ms moving window. This resulted in a time-dependent estimate of the matrix *A*. Eigenvalues of *A* closer to 0 indicate a faster decay, and eigenvalues near 1 would correspond to a slower decay. The method is also illustrated in Figures 7A-C and Figure S2E.

A z-test was applied to test for significant differences between the actual and shuffled results for the elements of the A matrices. The shuffled data were generated using the same matrix *X* but with randomized trial indexes across rows (*areas* × *variales*). The actual results with a mean outside the 99% confidence interval of the shuffled results showed a significant response.

#### One-dimensional task-variable-specific trajectory

We also carried out the targeted dimensionality reduction using condition-averaged pseudo-populations composed of 500 neurons recorded across sessions to study the population representation of gains and losses. We projected the condition-averaged population responses onto these axes:

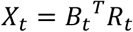

where *R*_*t*_ is condition-averaged population response at time *t*, which is of dimension *N*_*unit*_ × *N*_*condition*_. And *X*_*t*_ is the set of time-series vectors over all task variables and conditions, which is of dimension *N*_*f*_ × *N*_*condition*_.

The variability of trajectories across different conditions was estimated using a bootstrap procedure. According to the null hypothesis, we generated data with no differences among the conditions by sampling with replacement from the combined set of all conditions. We then calculated the sum of the standard error of each condition from the mean of all conditions for each bootstrap condition set. We did this 1000 times. It gave us 1000 different sampled standard errors from the null distribution. We then compared the standard error of the actual data to the standard errors in the null distribution. If the actual standard error is in the 99% confidence interval of the null distribution, the trajectories significantly differ among the conditions (i.e., p < 0.01).

### QUANTIFICATION AND STATISTICAL ANALYSIS

Unless otherwise indicated, all data were presented as means ± SEM (standard error of the mean). The statistical analyses performed were indicated in the main text and detailed in STAR Methods. Statistical comparisons were analyzed in MATLAB (Mathworks), including parametric, non-parametric, and permutation-based statistics, as detailed in STAR Methods. Figures were prepared with MATLAB.

