## Supplementary Information for "Ventral frontostriatal circuitry mediates the computation of reinforcement from symbolic gains and losses"

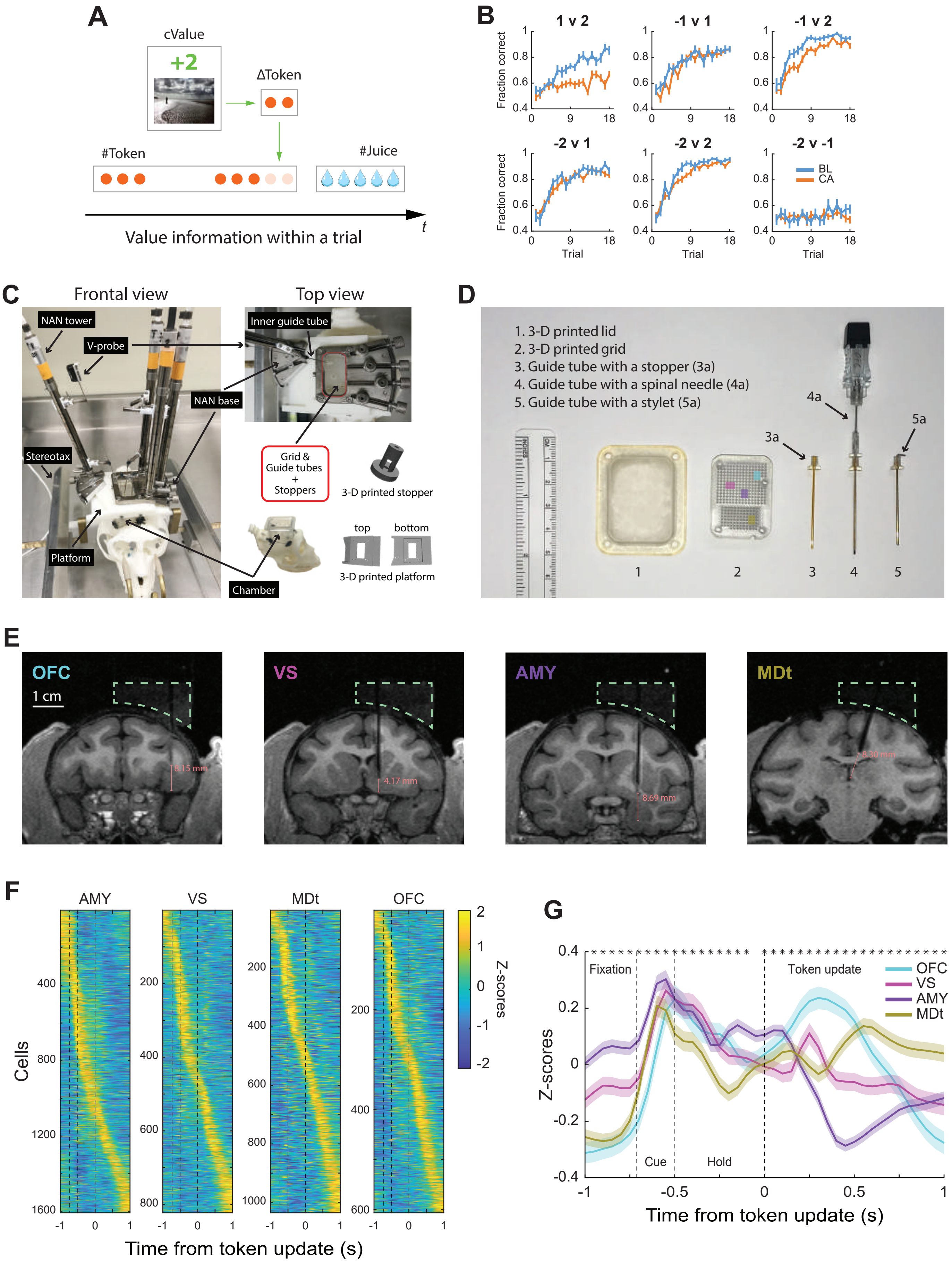
A
cValue
+2
ΔToken
#Token
#Juice
Value information within a trial
t
B
1 v 2
-1 v 1
-1 v 2
-2 v 1
-2 v 2
-2 v -1
Fraction correct
Trial
BL
CA
C
Frontal view
Top view
NAN tower
V-probe
Inner guide tube
NAN base
Stereotax
Platform
Chamber
Grid & Guide tubes + Stoppers
3-D printed stopper
top
bottom
3-D printed platform
D
1. 3-D printed lid
2. 3-D printed grid
3. Guide tube with a stopper (3a)
4. Guide tube with a spinal needle (4a)
5. Guide tube with a stylet (5a)
E
OFC
1 cm
VS
AMY
MDt
F
AMY
VS
MDt
OFC
Cells
Z-scores
Time from token update (s)
G
Fixation
Token update
Cue
Hold
OFC
VS
AMY
MDt
Z-scores
Time from token update (s)

Figure S1

### SUPPLEMENTAL FIGURE LEGENDS

#### Figure S1. Behavior, recorded areas, and neural activity, related to Figures 1 and 2.

(A) Different forms of value in the task. The primary reward was 1-12 drops of apple juice (#juice). The accumulated tokens were cashed out for juice every four to six trials. So, the reward signal was stored in the number of tokens (#token) for most trials. In each trial, the value update information was first carried by the images (cValue) and then transferred to the change of tokens (Δtoken).

(B) The choice behaviors of monkeys BL and CA.

(C) Recoding setup on a 3D-printed skull in a stereotaxic frame. Only one probe is shown.

(D) Components of the recording chamber. 1) 3-D printed lid. 2) 3-D printed grid, with vertical holes for the OFC, VS, and AMY (indicated by the colored squares), and holes with a 16° angel for the MDt; 3) Guide tube with a 3-D printed stopper glued at the top. 4) Guide tube with a spinal needle inside. This setup was used when inserting the guide tubes into the brain. The spinal needle penetrated the dura. 5) Guide tube with a stylet inside. This setup was used after the surgery. The stylets were about 0.5 mm longer than the guide tube protrusion to avoid clogging. They were removed during recording to allow probes to enter the guide tubes. All 3-D printed components were printed using biocompatible materials. All processes regarding guide tubes were carried out using sterile procedures.

(E) A coronal section view of the chamber and the shadow of a guide tube for each area under MRI. Orange lines indicate the longest distance the probes could advance from the tip of the guide tubes.

(F-G) Z-scored neural activity. (F) Heatmaps of the activity of all the neurons recorded from the AMY (n = 1607), VS (n = 829), MDt (n = 1035), and OFC (n = 606). Firing rates were z-scored based on activity during the presented period. Warmer colors indicate higher firing rates. Neurons were aligned to the token update and rank-ordered by the latency of the response peak. One row represents one neuron. (G) Mean z-scored activities of each area. Shaded zones represent mean ± SEM. The black asterisks indicate a significant difference among the four areas (one-way ANOVA, $p < 0.01$). Vertical black dashed lines represent the onset of images, the start of holding on the chosen image, and token update, which split the time course into four epochs: fixation, cue, hold, and token update.

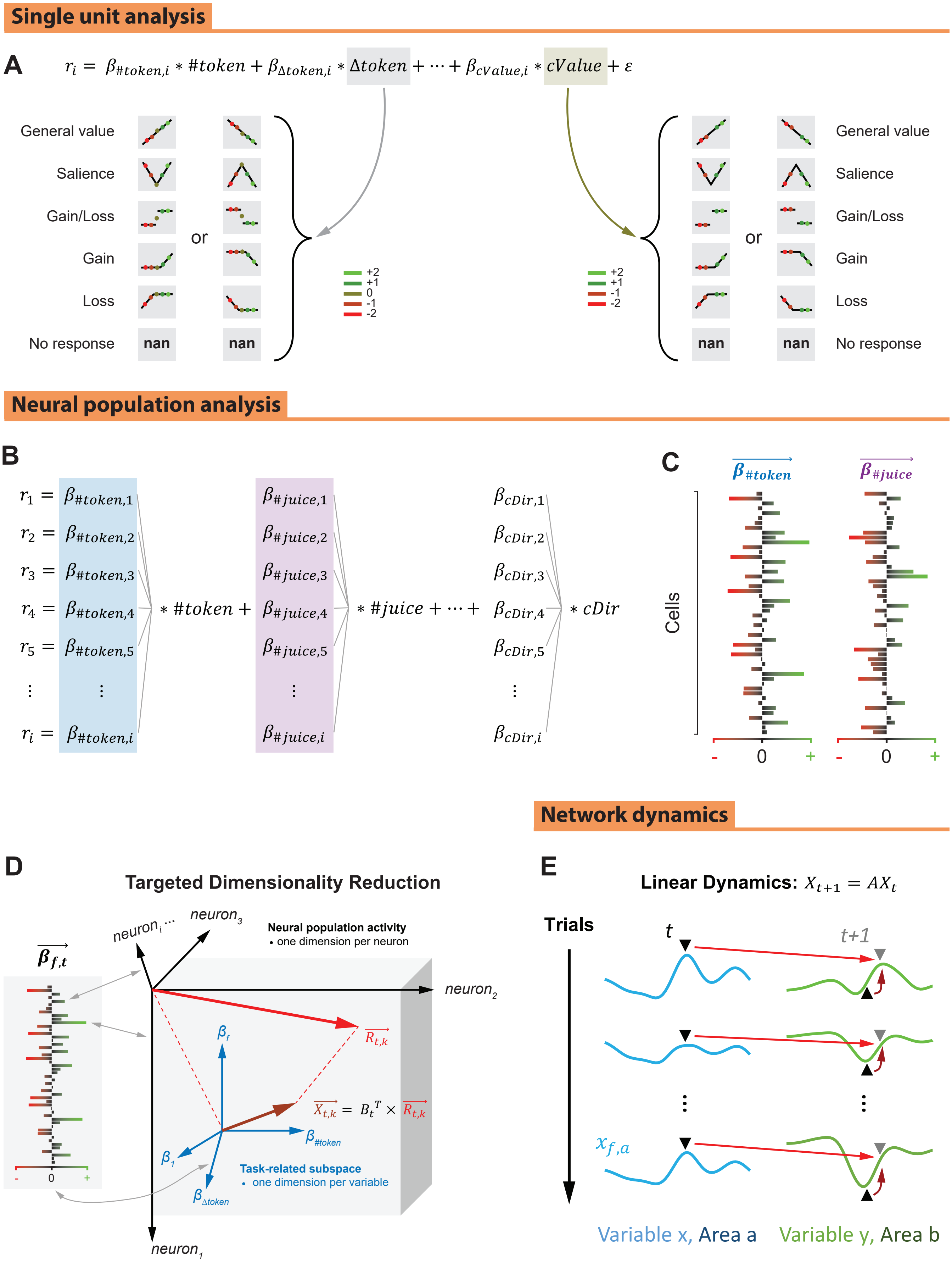
Single unit analysis
A
$r_i = \beta_{\#token,i} * \#token + \beta_{\Delta token,i} * \Delta token + \cdots + \beta_{cValue,i} * cValue + \varepsilon$
General value
Salience
Gain/Loss
Gain
Loss
No response
or
nan
+2
+1
0
-1
-2
Neural population analysis
B
$r_1 = \beta_{\#token,1}$
$r_2 = \beta_{\#token,2}$
$r_3 = \beta_{\#token,3}$
$r_4 = \beta_{\#token,4}$
$r_5 = \beta_{\#token,5}$
$r_i = \beta_{\#token,i}$
$* \#token +$
$\beta_{\#juice,1}$ … $\beta_{\#juice,i}$
$* \#juice + \cdots +$
$\beta_{cDir,1}$ … $\beta_{cDir,i}$
$* cDir$
C
$\overrightarrow{\beta_{\#token}}$
$\overrightarrow{\beta_{\#juice}}$
Cells
- 0 +
Network dynamics
D
Targeted Dimensionality Reduction
$\overrightarrow{\beta_{f,t}}$
$neuron_i$ …
$neuron_3$
Neural population activity
• one dimension per neuron
$neuron_2$
$neuron_1$
$\beta_f$
$\overrightarrow{R_{t,k}}$
$\overrightarrow{X_{t,k}} = B_t^T \times \overrightarrow{R_{t,k}}$
$\beta_1$
$\beta_{\#token}$
$\beta_{\Delta token}$
Task-related subspace
• one dimension per variable
E
Linear Dynamics: $X_{t+1} = AX_t$
Trials
t
t+1
$x_{f,a}$
Variable x, Area a
Variable y, Area b

Figure S2

#### Figure S2. Systematic procedures for analyzing neural population data at single-cell, population, and network levels, related to Figures 3-7.

(A) Classifying neurons by replacing independent variables (in this case, Δtoken and cValue) with several potential combinations in multivariate linear regression models. A neuron was classified into the categories as the combination included in the best-fitting model. Nan means removing the variable from the model.

(B-C) Approach to calculate population coding similarity. (B) The regression coefficient vector, $\overrightarrow{\beta_f}$, of a neural population represents the weights of each neuron in the population encoding one task variable $f$. It indicates the relationship between neural population activity and the task variable. (C) The cross-correlation between two regression coefficient vectors (simulation of 50 neurons) indicates how similar a neural population encoded the two task variables.

(D) Targeted dimensionality reduction. Task-variable-specific latent variable at time $t$ on trial $k$, $\overrightarrow{X_{t,k}}$ (one vector in the matrix $X_t$), was calculated by projecting single-trial population activity $\overrightarrow{R_{t,k}}$ (one vector in the matrix $R_t$) into the subspace generated by axes $B_t$, which consisted of $\overrightarrow{\beta_{f,t}}$. Each coefficient vector $\overrightarrow{\beta_{f,t}}$ maps the relationship between all the axes in the high-dimensional space (one $\beta$ for one axis) and the task variable $f$ axis at time $t$ in the low-dimensional subspace. We also carried out the same approach using condition-averaged population response, but not at the single trial level, analyzing the population representation of gains and losses. See also Methods.

(E) Linear dynamics. Matrix $A$ measures the within-area/variable interactions (dark red arrows) and across-area/variable interactions (light red arrows) at each point in time. The one-dimensional single-trial task-variable-specific latent variable $x_{f,a}$ consisted of the projections of $\overrightarrow{X_{t,k}}$ from area $a$ on the axis $\beta_f$ across the time course of the trial.

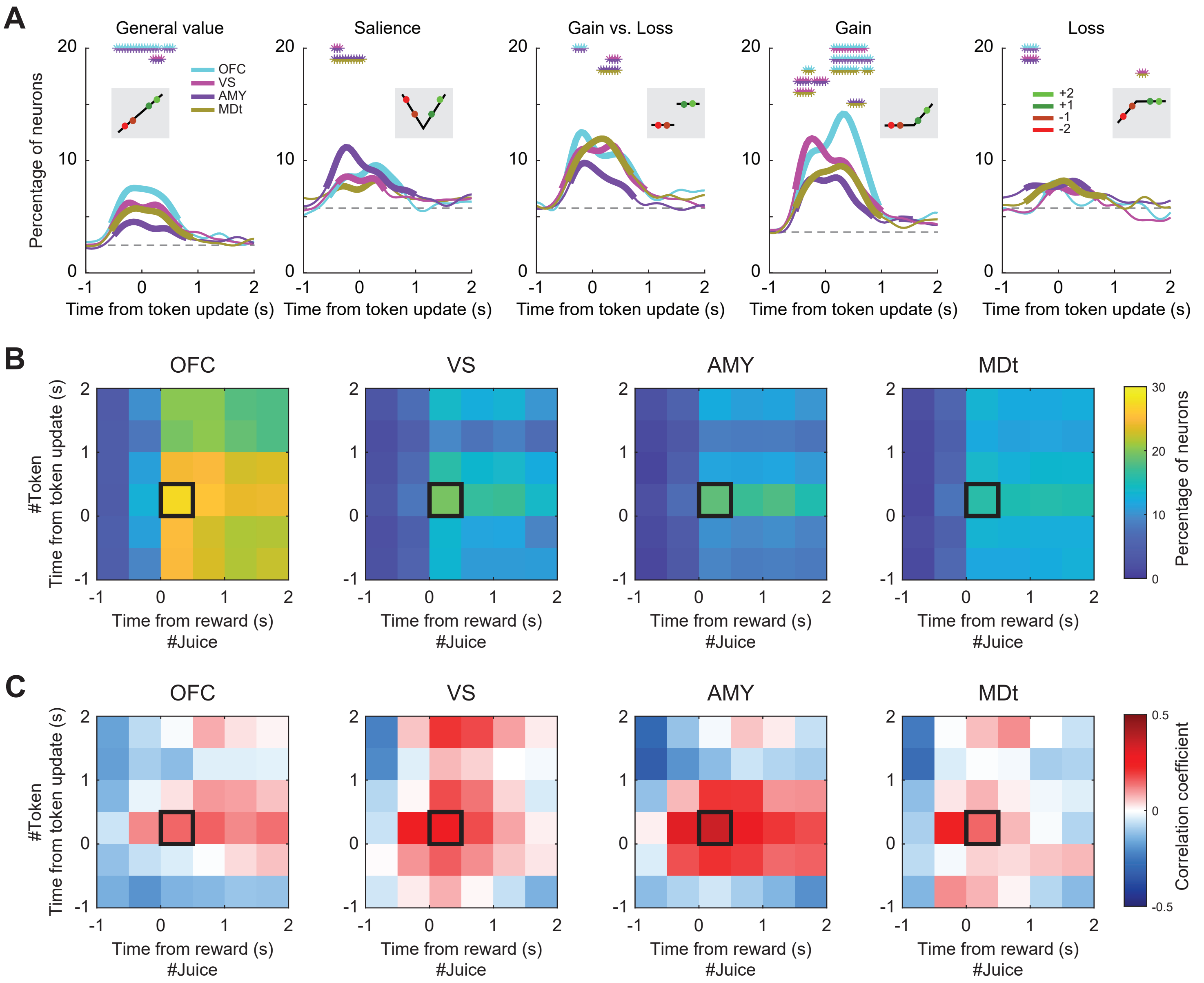
A
General value
Salience
Gain vs. Loss
Gain
Loss
Percentage of neurons
OFC
VS
AMY
MDt
+2
+1
-1
-2
Time from token update (s)
B
OFC
VS
AMY
MDt
#Token
Time from token update (s)
Time from reward (s)
#Juice
Percentage of neurons
C
Correlation coefficient

Figure S3

#### Figure S3. Coding of chosen value, tokens, and the number of juice rewards, related to Figures 3 and 4.

(A) Percentage of neurons in each area encoding the *a priori* value of chosen images (cValue).

(B) Percentage of neurons encoding #token and #juice. The x-axis represents #juice, and the y-axis represents #token. Black squares indicate the time windows used for the bar plots in Figures 4H-I.

(C) Encoding similarity between #token and #juice in each area.

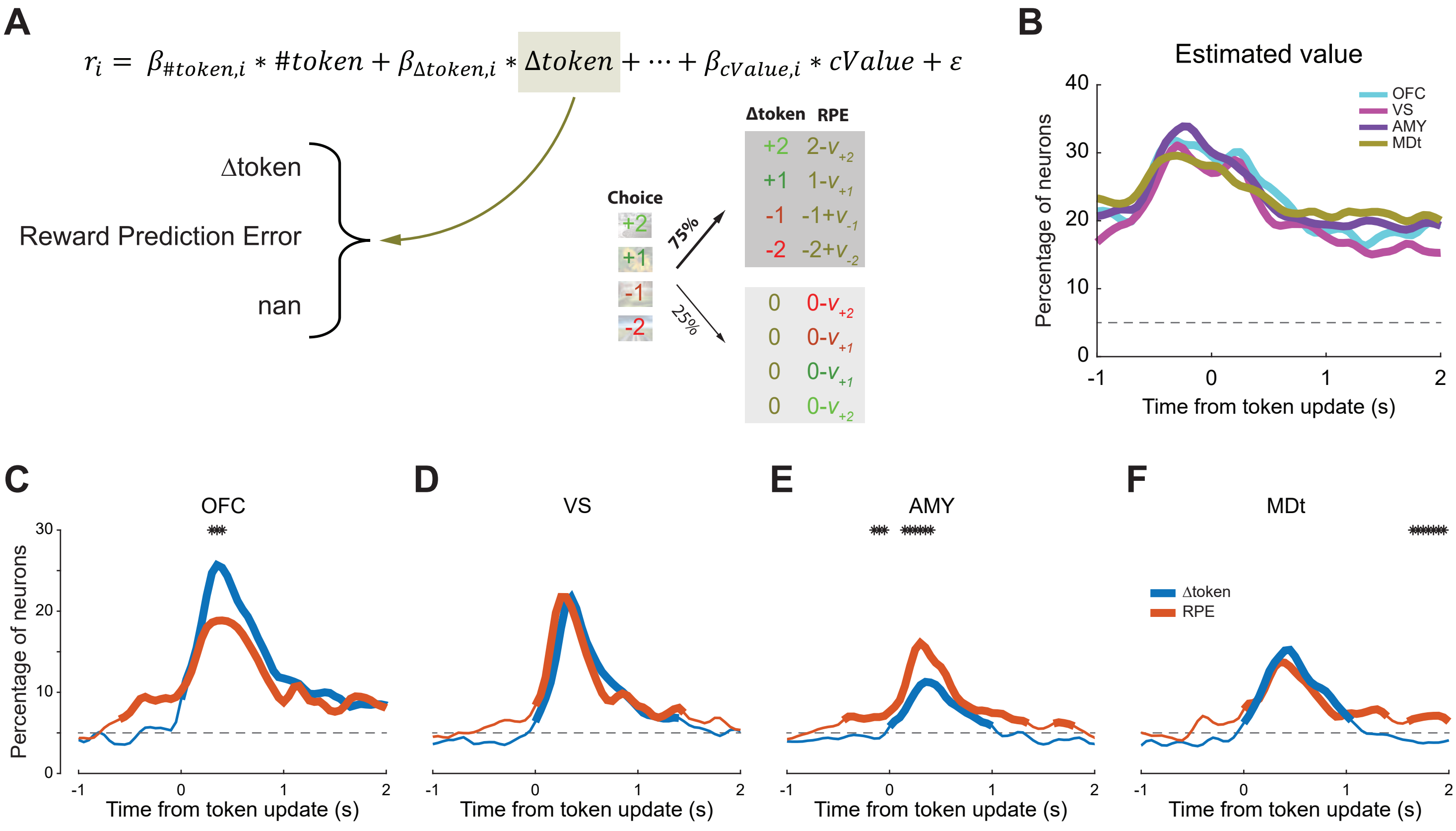

A
$r_i = \beta_{\#token,i} * \#token + \beta_{\Delta token,i} * \Delta token + \cdots + \beta_{cValue,i} * cValue + \varepsilon$
Δtoken
Reward Prediction Error
nan
Choice
+2
+1
-1
-2
75%
25%
Δtoken RPE
+2 2-$v_{+2}$
+1 1-$v_{+1}$
-1 -1+$v_{-1}$
-2 -2+$v_{-2}$
0 0-$v_{+2}$
0 0-$v_{+1}$
0 0-$v_{+1}$
0 0-$v_{+2}$
B
Estimated value
OFC
VS
AMY
MDt
Percentage of neurons
Time from token update (s)
C
OFC
D
VS
E
AMY
F
MDt
Δtoken
RPE
Percentage of neurons
Time from token update (s)


Figure S4

#### Figure S4. Value updates in the ventral striatum and orbitofrontal cortex, related to Figure 5.

(A) Replacing the independent variable Δtoken with potential combinations (Δtoken, RPE, or nan). A neuron was classified into the category that the best-fitting model indicated.

(B) Percentage of neurons in each area encoding the value of chosen images, $v_h$, estimated by the RW model. Dashed horizontal lines represent the chance level. Thick lines indicate a significant difference between the corresponding area and chance level (binomial test, $p < 0.01$).

(C-F) Percentage of neurons in each area encoding Δtoken and RPE. Dashed horizontal lines represent chance levels. Thick lines indicate a significant difference between the corresponding area and chance level (binomial test, $p < 0.05$). The black asterisks indicate a significant difference between the two categories (chi-square test, $p < 0.01$).

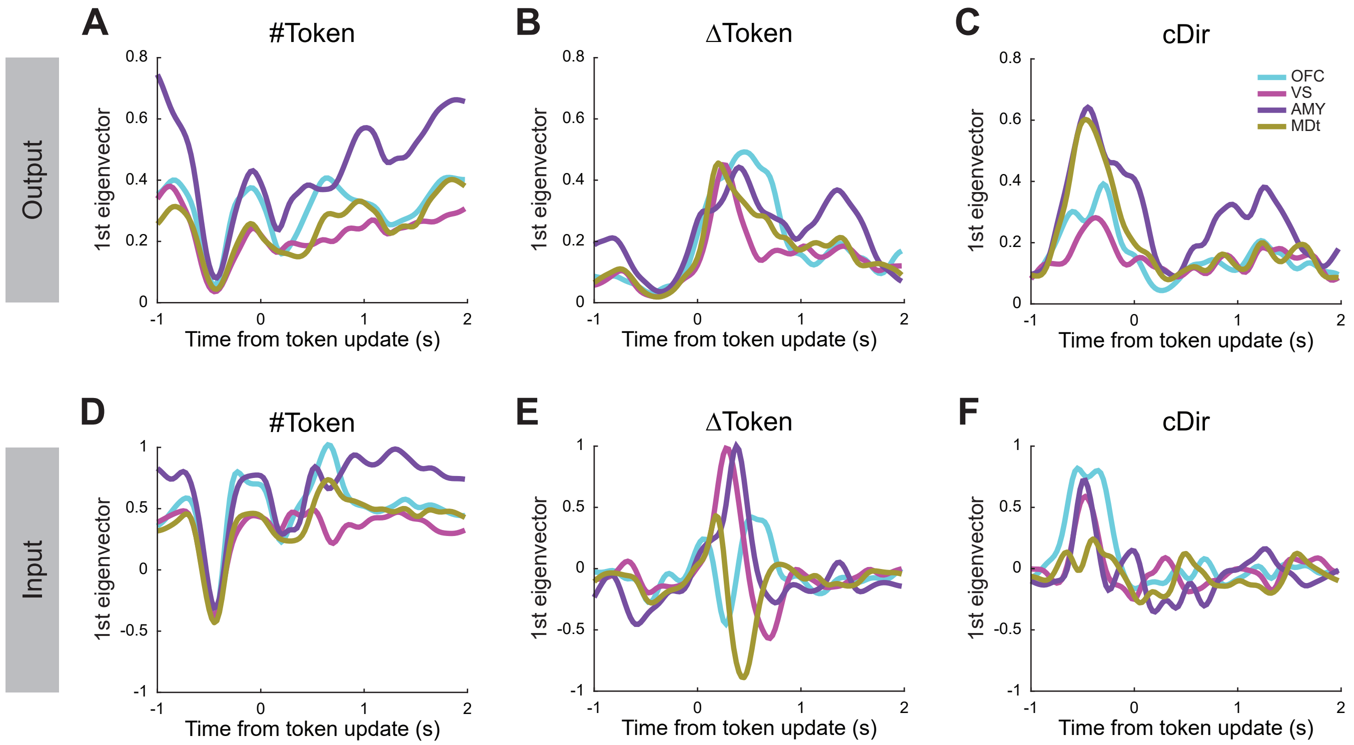
Output
Input
A
#Token
B
ΔToken
C
cDir
D
#Token
E
ΔToken
F
cDir
1st eigenvector
Time from token update (s)
OFC
VS
AMY
MDt

Figure S5

#### Figure S5. Eigenvectors of the matrix A, related to Figure 7.

(A-C) The columns of eigenvectors capture the information flowing from the source areas and task variables.

(D-F) The rows of inversed eigenvectors capture the information flowing into the source areas and task variables.

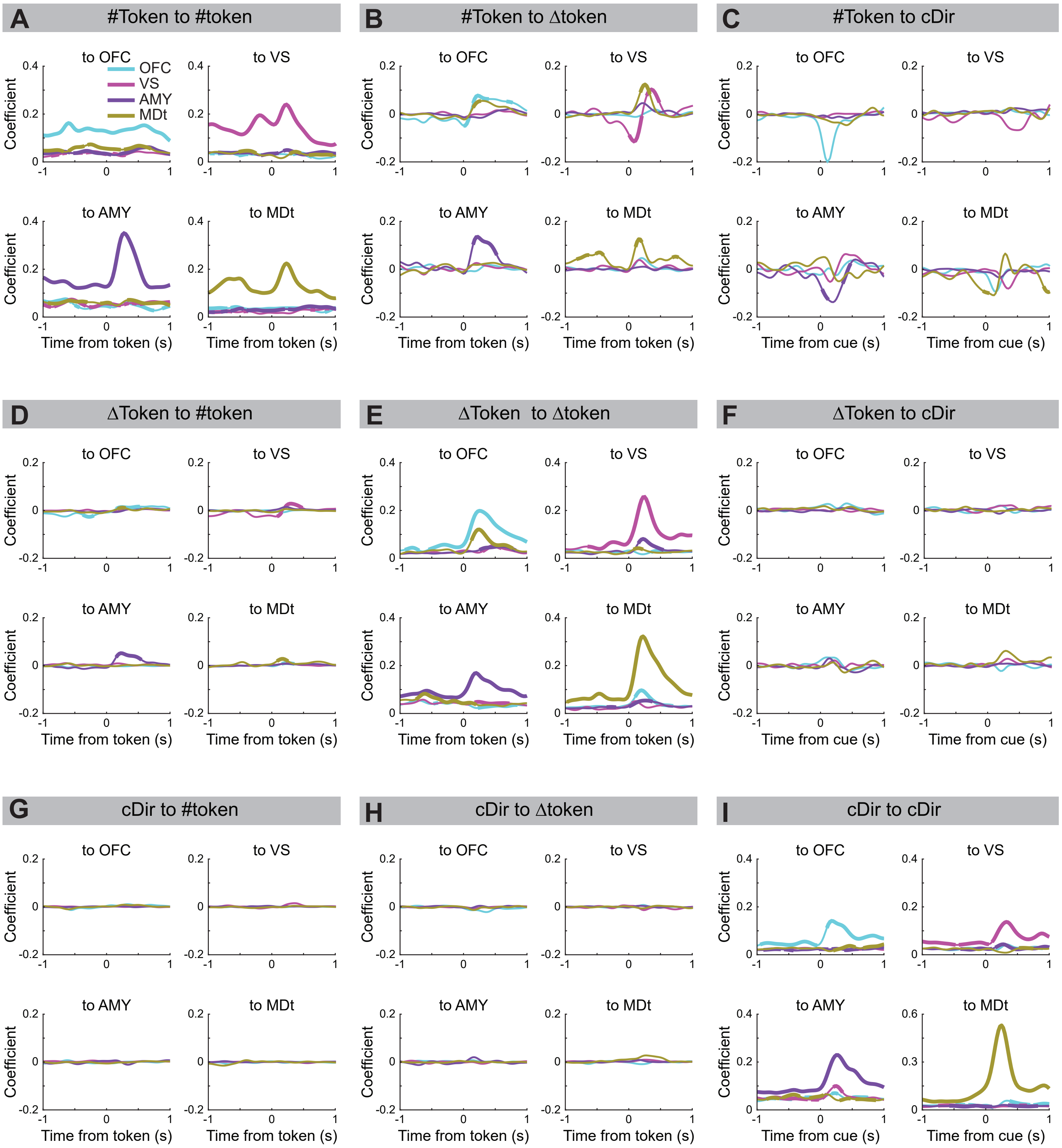
A
#Token to #token
B
#Token to Δtoken
C
#Token to cDir
D
ΔToken to #token
E
ΔToken to Δtoken
F
ΔToken to cDir
G
cDir to #token
H
cDir to Δtoken
I
cDir to cDir
to OFC
to VS
to AMY
to MDt
OFC
VS
AMY
MDt
Coefficient
Time from token (s)
Time from cue (s)

Figure S6

#### Figure S6. Flow of token and direction information in the ventral network, related to Figure 7.

(A-I) The flow of token and chosen direction information across areas and task variables. All combinations among the three task variables are shown.
